# Decoding the Dual Recognition Mechanism of Glucocorticoid Receptor for DNA and RNA: Sequence vs. Shape

**DOI:** 10.1101/2023.03.15.532803

**Authors:** Hörberg Johanna, Anna Reymer

## Abstract

Transcription factors (TFs) regulate eukaryotic transcription through selecting DNA-binding, can also specifically interact with RNA, which may present another layer of transcriptional control. The mechanisms of the TFs-DNA recognition are often well-characterised, while the details of TFs-RNA complexation are less understood. Here we investigate the dual recognition mechanism of the glucocorticoid receptor (GR), which interacts with similar affinities with consensus DNA and diverse RNA hairpin motifs but discriminates against uniform dsRNA. Using atomic molecular dynamics simulations, we demonstrate that the GR binding to nucleic acids requires a wide and shallow groove pocket. The protein effectively moulds its binding site within DNA major groove, which enables base-specific interactions. Contrary, the GR binding has little effect on the grooves geometry of RNA systems, most notably in uniform dsRNA. Instead, a hairpin motif in RNA yields a wide and shallow major groove pocket, allowing the protein to anchor itself through nonspecific electrostatic contacts with RNA backbone. Addition of a bulge increases RNA hairpin flexibility, which leads to a greater number of GR-RNA contacts and, thus, higher affinity. Thus, the combination of structural motifs defines the GR-RNA selective binding: a recognition mechanism, which may be shared by other zinc finger TFs

## INTRODUCTION

Transcription factor proteins, via selective binding to DNA, regulate genetic transcription (1). Multiple experiments suggest that transcription factors (TFs) also specifically interact with RNA, which may present an additional layer of transcription regulation (2–4). RNA molecules are constantly present at the actively transcribed sites (5–9). As our understanding of the role of TFs-RNA interactions still evolves, the available data suggest that TF-RNA interactions can facilitate the association of TFs with the chromatin fibre and the search for their corresponding DNA binding sites. Examples include, nuclear TF Y (NF-YA) interactions with PANDA long noncoding RNA (10), which titrates the TF from the target genes to reduce their expression; Yin Yang 1 TF (YY1) interactions with nascent transcripts to retain its presence at actively transcribed promoters (11); heat shock factor 1 (HSF1) that employs RNA as a scaffold for the TFs assembly on DNA (12). Furthermore, a recent study by Oksuz et al. reports that nearly half (48%) of TFs identified in human K562 cells appear to interact with RNA (13). As TF proteins do not possess the characteristic structural domains of well-studied RNA binding proteins (14, 15), little is known on the structural basis of RNA-TF associations.

Here we aim to provide atomic level insights into what drives the TF-RNA association, using as a model system the glucocorticoid receptor that can interact with RNA (4, 16–18). Glucocorticoid receptor (GR) is a ubiquitous TF, which regulates the transcription of thousands of genes associated with development, immune responses, metabolism, inflammation, apoptosis, etc (19–21). To regulate a transcriptional response, the GR needs a glucocorticoid binding, which induces the factor translocation from the cytoplasm to the nucleus to enable interactions with DNA. GR is widely dysregulated in disease, and thus is targeted by synthetic glucocorticoids to combat a range of autoimmune disorders and many cancers, as a part of combinatorial therapy (22–24). GR has a modular structure (25), including two well-characterised domains: a ligand-binding domain and a DNA-binding domain. The GR DNA-binding domain (DBD) consists of two Zn-fingers of Cys4-type, and associates on DNA in several configurations that lead to distinct transcriptional responses (21, 26–29). The GR dimer-DNA association, in a head-to-head orientation, leads to the transcription induction. The GR monomer-DNA association, collaboratively or tethered to other TFs, e.g., from AP-1 or COUP families, leads to the transcription repression. The GR-DBD-DNA binding is highly sequence-specific. The protein recognises the AGAACA sequence, known as the glucocorticoid response element (GRE).

The GR interactions with RNA are much less explored. Several reports describe GR associations with tRNA, mRNA, and Gas5 long noncoding RNA (lncRNA) (16–18). It has been proposed that GR binds the GRE-like elements within RNA molecules that act as molecular decoys for the GR-DBD, but details on the mechanism and selectivity remain elusive. An NMR study by Parsonnet et al. (4), focusing on the mechanism of Gas5 lncRNA-GR interactions *in vitro*, reported intriguing results. Firstly, GR-DBD interacts with RNA as a monomer and adopts a distinct RNA-bound state, different from the DNA-bound and free states. Secondly, the protein recognises a range of RNA hairpin motifs, both synthetic and biologically derived, but discriminates against uniform dsRNA. And finally, the experiments suggested that GR recognises RNA in a structure-specific and not in sequence-specific manner, as is in the DNA case.

Motivated by these experimental insights, we want to further explore the mechanistic details of the GR-RNA recognition, using all-atom molecular dynamics simulations. As target systems for the GR binding, we employ several RNA structures: a fully paired hairpin, a hairpin with a bulge, and a uniform dsRNA; and compare those to the GR interactions with its native DNA target. All studied nucleic acids contain the GRE-site. We observe that the presence of a hairpin and a bulge in an RNA molecule creates the most suitable pocket for the GR binding, which results in a similar interaction energy as is in the DNA-GR monomer case. However, contrary to GRE-DNA-GR recognition, which requires the formation of several specific contacts, the GR binding to RNA is secured by nonspecific electrostatic interactions of long-chain positively charged residues to RNA backbone. Furthermore, our data suggest that it is rather the 3D structural motif of an RNA molecule rather than its sequence that determines the stable binding by GR.

## METHODS

### Systems Preparation

We simulate four different nucleic acids systems where the glucocorticoid receptor (GR) interacts with DNA (CCAGAACAGAGTGTTCTGA), dsRNA (GGAGAACAAAAUGUUCUUU), fully paired RNA hairpin “GRE_RNA” (GGCAAAAUGUUCUUUCGAGAACAUUUUGCC) and bulge-containing hairpin “Gas5” (GGCCCAGUGGUCUUUGUAGACUGCCUGAUGGCC) (Figure 1), and their unbound counterparts (4). We derive the 3D structures for the GRE_RNA and Gas5_RNA hairpins through secondary structure prediction with RNAFold webserver (30, 31) (http://rna.tbi.univie.ac.at/cgi-bin/RNAWebSuite/RNAfold.cgi) followed by 3D structure prediction with 3dRNA webserver (32, 33) (https://bio.tools/3dRNA). We create the A-form dsRNA structure in USCF Chimera (34). To derive GR-RNA complexes, we perform the macromolecular docking in HDOCK (35), where GR (PDB ID: 6CFN (36)) is assigned as the receptor and the RNA molecule – as the ligand. The modelling of Gas5_RNA provides a larger structural variation compared to GRE_RNA (Figure S1), thus we use two models of Gas5 termed, model 1 and model 2, for the docking. In addition, we exclude helix 4 of GR (Figure 1) during the docking, to obtain a similar GR-RNA interacting pose as is in the GR-DNA complex as helix 4 might provide a steric hindrance. We employ this docking setup, as the NMR experiments (4) imply that GR binds RNA in a similar fashion as DNA. The docking generates the GR-RNA complexes matching this criterion among the top-ten scored decoys (Figure S2). Experiments also report that GR binds DNA as a dimer, whereas RNA as a monomer. The dimeric GR-DNA complex is built in USCF Chimera from the GR-DNA complex crystal structure (PDB ID: 5CBX)(37), where each GR monomer was exchanged for the helix4 containing GR crystal structure (PDB ID: 6CFN).

**Figure 1.**
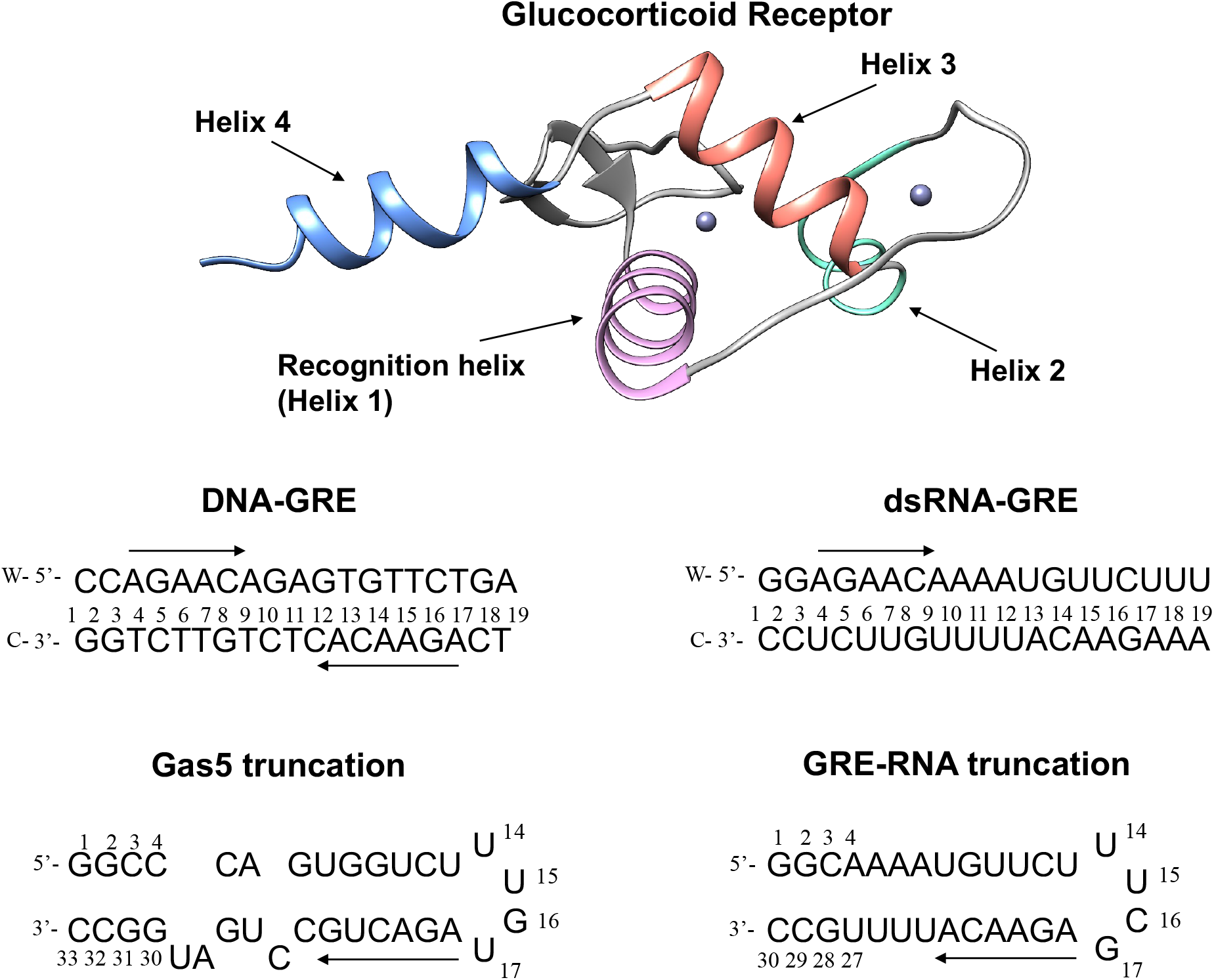
The 3D structure of the glucocorticoid receptor (PDB ID: 6CFN (36)), with indicated recognition helix, central for the selective DNA binding; and helix 4, implicated in binding of RNA. Below are the 2D structures of the studied nucleic acids systems with their nucleotide sequences. The direction of the GR-response element (GRE) is highlighted with an arrow.

### Molecular Dynamics Simulation

We perform all molecular dynamics simulations with the GROMACS MD engine version 2020.5,(38) using amber14SB (39), parmbsc1 (40) and parmbsc0+chiOL3 (41, 42) forcefields for the protein, DNA and RNA, respectively. GR contains two ZnCys_4_clusters, which we treat with the ZAFF parameters (43). We solvate DNA and GR-DNA systems with 15Å SPC/E (44) water in cubic boxes and neutralise with K^+^. We add the K^+^ and Cl^−^ ions to reach a physiological concentration of 150mM KCl. We employ a different setup for all RNA simulated systems, which we solvate with 15Å TIP3P water (44) in cubic boxes and neutralise with Na^+^. We add the Na^+^ and Cl^−^ ions to reach a concentration of 100mM NaCl. We treat the monovalent ions with the Joung-Cheatham parameters (45). We proceed with energy minimization with 5000 steps of steepest decent, followed by 500 ps equilibration-runs with weak position restraints on heavy atoms of the solute (1000 kJ/mol) in the NVT and NPT ensembles, adjusting temperature and pressure to 300 K and 1 atm, (46, 47) in periodic boundary conditions. For all RNA systems, both unbound and GR-bound, we perform an additional equilibration run with position restraints on heavy atoms of the solute (1000 kJ/mol) in the NPT ensemble for 20ns (48). Releasing the restraints, for each of the systems we carry out 600ns MD simulations at constant pressure and temperature (1 atm and 300 K).

### Trajectory Analysis

For each of the generated MD trajectories, we discard the first 100 ns as equilibration. To analyse the protein-nucleic acids contacts, we use the CPPTRAJ (50) program from AMBERTOOLS 16 software package. We distinguish between intermolecular specific (hydrogen bonding and hydrophobic contacts formed between protein side chains and nucleic acids bases) and nonspecific contacts (that involve interactions with either DNA/RNA or protein backbones). We consider only the contacts that are present for longer than 10% of the trajectory, see previous publications for the details (49, 53). Subsequently, we apply Curves+ and Canal programs (51) to derive the helical parameters, backbone torsional angles and groove geometry parameters of the nucleic acid systems for each trajectory snapshot extracted every ps. For the RNA hairpin systems, we perform the analysis for the fully paired GRE-site region. We use the GROMACS energy tool to calculate the protein-DNA/RNA interaction energies, which include short range electrostatic and Lennard-Jones interactions. To separate interaction energies into specific and nonspecific, we calculate interactions for several atom groups that correspond to DNA/RNA bases, protein side chains, and molecule backbone. We employ the GROMACS “covar” and “anaeig” tools to calculate the configurational entropies, using two subsets of atoms: the backbone “P” atoms of the entire DNA/RNA sequences and of only the GR-response element. From the obtained eigenvectors, we calculate the configurational entropies at 300 K using the Schlitter’s formula (52). To derive standard deviations, we calculate the configurational entropies for the entire 500 ns trajectory and for each consecutive 100ns windows of the trajectory. We perform the principal component analysis (PCA) for every MD trajectory using the GROMAS “covar” tool. We perform PCA for the unbound and the GR-bound systems. For the GR-bound systems, we use three different PCA protocols: (1) for the heavy atoms of the protein-DNA/RNA complexes with the superposition on the nucleic acids; (2) for the heavy atoms of DNA/RNA with the superposition on the nucleic acids; and (3) for the heavy atoms of the protein-DNA/RNA complexes with the superposition on the entire systems.

### Additional Information

We use MatLab software for the post-processing and plotting of all data and USCF Chimera (34) for all molecular graphics.

## RESULTS AND DISCUSSION

*In vitro* binding studies demonstrated that the GR-DBD (DNA-binding domain) associates with high affinity with Gas5 lncRNA (16, 18), recognising RNA hairpin motifs within and discriminating against dsRNA (4). To rationalize the experimental observations and gain further insights into the mechanism of GR association with RNA, we conduct a computational study. As target systems for the GR binding, we employ B-DNA containing the protein native target sequence and several RNA structures: a fully paired hairpin, a hairpin with a bulge, and uniform dsRNA (Figure 1). The DNA molecule, 19 b.p. in length, contains two GRE-sites (AGAACA) separated by a three b.p. spacer. Same sequence composition is also selected for the dsRNA molecule, as it is designed to mimic the DNA system. The fully paired RNA hairpin (later ‘GRE_RNA’), 30 nucleotides in length, contains the GRE-motif followed by the terminal loop UUCG (5’->3’). The RNA hairpin with a bulge (later ‘Gas5_RNA’), 33 nucleotides in length, is derived from noncoding Gas5 RNA, contains the GRE-like motif (AGACUG) followed by the same terminal loop as is in GRE_RNA.

We derive the models for Gas5_RNA and GRE_RNA through the secondary structure prediction in RNAFold (30, 31) followed by the 3D structure prediction in 3dRNA (32, 33). We build the A-form dsRNA molecule using USCF Chimera (34). To derive the GR-RNA complexes we perform the protein-nucleic acids docking in HDOCK (35), specifying GR (PDB ID: 6CFN (36)) as a “receptor” and RNA as a “ligand”. The modelling of the 3D structure of Gas5_RNA provides larger structural variations in the hairpin region compared to GRE_RNA (Figure S1). Thus, we derive the GR-Gas5_RNA complexes for the two highest scored Gas5_RNA models (later termed model 1 and model 2). The NMR experiments (4) suggest that GR-DBD binds to RNA in a similar fashion as to DNA. However, the stoichiometry of the binding differs: GR binds DNA as a dimer, whereas RNA as a monomer. The docking in HDOCK results in GR-Gas5_RNA, GR-GRE_RNA, and GR-dsRNA complexes matching these criteria among the top-ten scored decoys. We build the GR dimer-DNA complex in USCF Chimera from the crystal structure of the complex (PDB ID: 5CBX (37)) where we exchange each GR monomer with the GR crystal structure (PDB ID: 6CFN) that includes helix 4, which was shown to be important for the GR-RNA association. We subsequently subject the derived five GR-nucleic acids complexes as well as all unbound RNA and DNA systems to 600 ns all-atomistic molecular dynamics (MD) simulations.

We first compare the trajectory-averaged structures for the bound and unbound DNA and RNA molecules, to see if the binding of GR contributes to any significant conformational changes in nucleic acids (Figure 2A). DNA, GRE_RNA and dsRNA exhibit no major conformational changes, illustrated by the RMSD values of ∼1 Å. Gas5_RNA shows greater conformational changes upon the binding of GR with the RMSD values of ∼3 Å, however, within the GRE-site, the changes are small, with the RMSD values of ∼1 Å. This suggests that the binding site within the nucleic acids systems is predefined by the 3D structure of a particular sequence.

**Figure 2:**
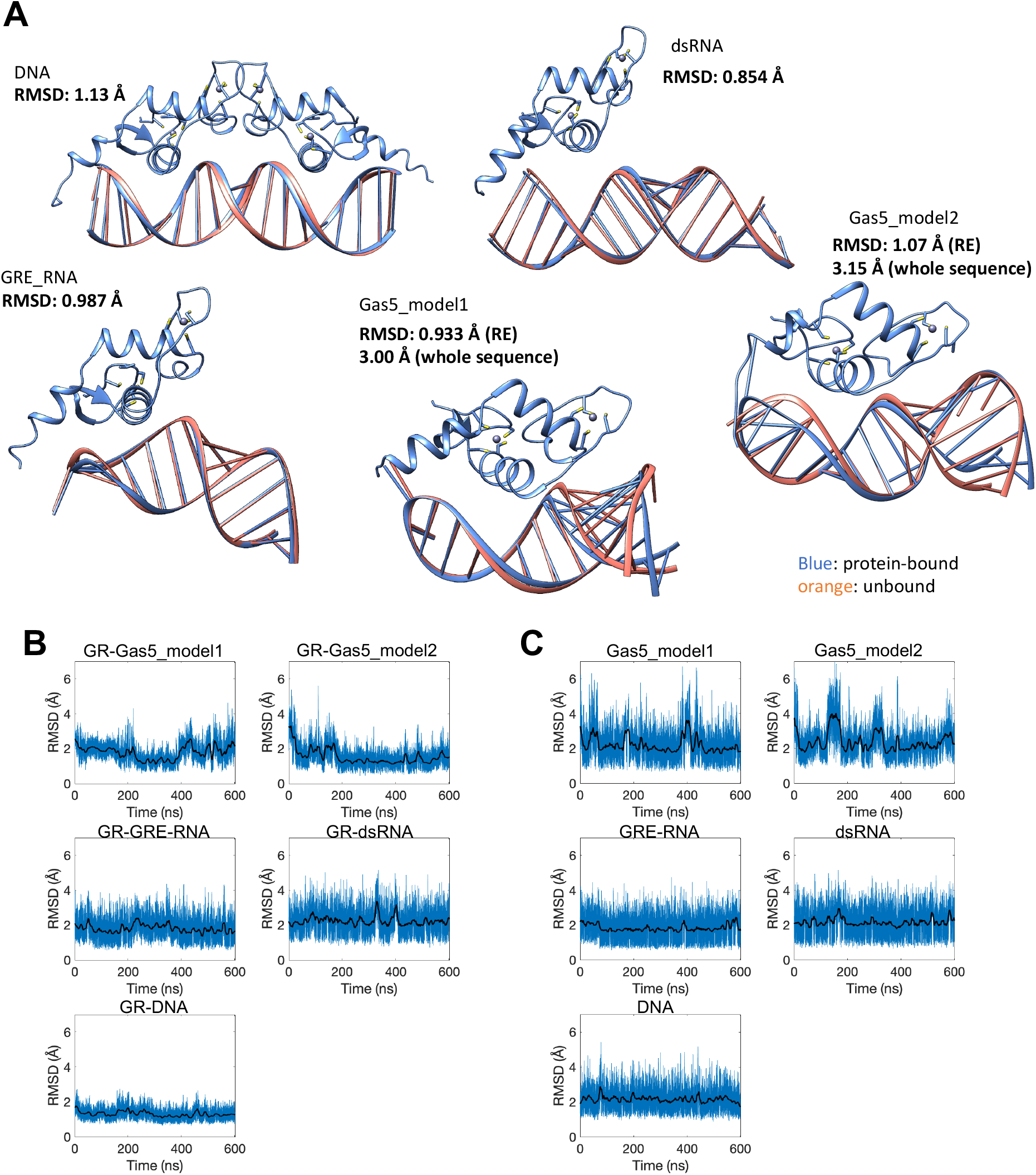
**A:** Comparison of the trajectory-averaged structures for the unbound and GR-bound DNA, dsRNA, GRE_RNA and Gas5_RNA systems. Evolution of the RMSD values along the trajectories for **B:** GR-bound-DNA/RNA and **C:** unbound DNA/RNA. The RMSD values are calculated for heavy atoms of DNA/RNA, with respect to the corresponding average structures.

We next analyse the evolution of the RMSD values of heavy atoms of DNA/RNA with respect to the average state along the MD trajectories. As seen in Figure 2B-C, the binding of GR to DNA and Gas5_RNA significantly reduces the RMSD values fluctuations compared to the unbound state. For GRE_RNA, the fluctuations of the RMSD values are smaller, whereas for dsRNA, the GR binding has no impact on the RMSD behaviour. The decreased fluctuations of the RMSD values, upon the protein binding, suggests a greater surface complementarity of the interacting molecules and the stability of the intermolecular contacts networks in the GR-DNA and GR-Gas5_RNA complexes.

### Protein–Nucleic Acids Contacts

We continue exploring the differences in the GR-DNA and GR-RNA binding mechanisms through the analysis of the protein-DNA/RNA contact networks. For this, we employ our dynamic contacts map approach (49, 53), where we follow the evolution of the number and strength of specific and nonspecific contacts (Figures S3-S8). By specific contacts we mean the contacts formed between atoms of nucleic acids bases and protein side chains, and by nonspecific contacts – the contacts that involve atoms of either molecule backbones.

In case of the GR–DNA binding, each GR monomer specifically interacts with the GRE-site (AGAACA) employing mainly the residues of the recognition helix: His432, Lys442, Val443, and Arg447 (Figure 3A). The intermolecular contact network is entangled, including specific hydrogen bonds and hydrophobic contacts as well as nonspecific electrostatic interactions (Figures 3A, 4). In detail, for GR monomer 1, His432 interacts with the outer CA step, including the A nucleotide of the response element (**C-A**GAACA). Lys442 interacts with the AG step (**AG**AACA); and Val443 and Arg447 form hydrophobic contacts with the GT step and the last T nucleotide, respectively, on the opposite DNA strand (AGAACA/**TGT**TCT). Arg447 also forms nonspecific electrostatic interactions with DNA backbone. For GR monomer 2, the residues exhibit nearly identical DNA-contacts as the monomer 1, with a few differences: His432 forms no specific contacts with the response element flanking sites while Lys442 exhibits stronger specific contacts. The alanine scanning experiments (4) provide the contribution order of the positively charged residues of the GR-DBD domain for the DNA binding: R447>K471>K480≈R479>K446. Based on our simulations, we conclude that R447 plays the major role for the GR-DNA binding and recognition: the residue interacts with the GRE-site trough specific and nonspecific contacts. The K471 and K446 residues interact with DNA backbone, while K480 and R479 may play a structural stability role as these are not observed to directly interact with DNA.

**Figure 3:**
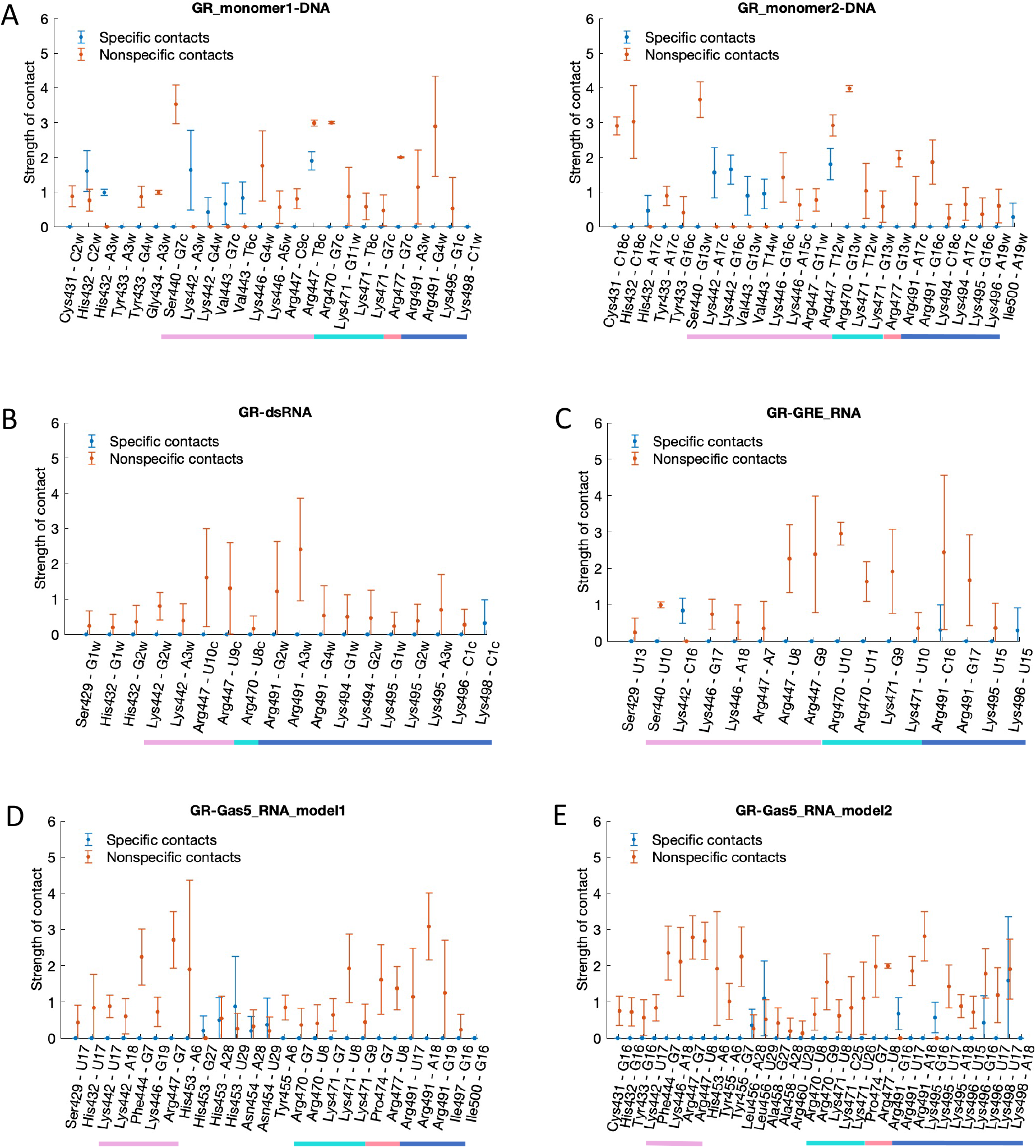
Average contacts strength with standard deviations for specific (blue) and nonspecific (orange) contacts for **A**. GR-DNA **B**. GR-dsRNA **C**. GR-GRE_RNA **D**. GR-Gas5_model1 **E**. GR-Gas5_model2. The different helices of GR-DBD are denoted with coloured lines: H1: pink, H2: aquamarine, H3: coral, H4: blue (see also Figure 1).

**Figure 4.**
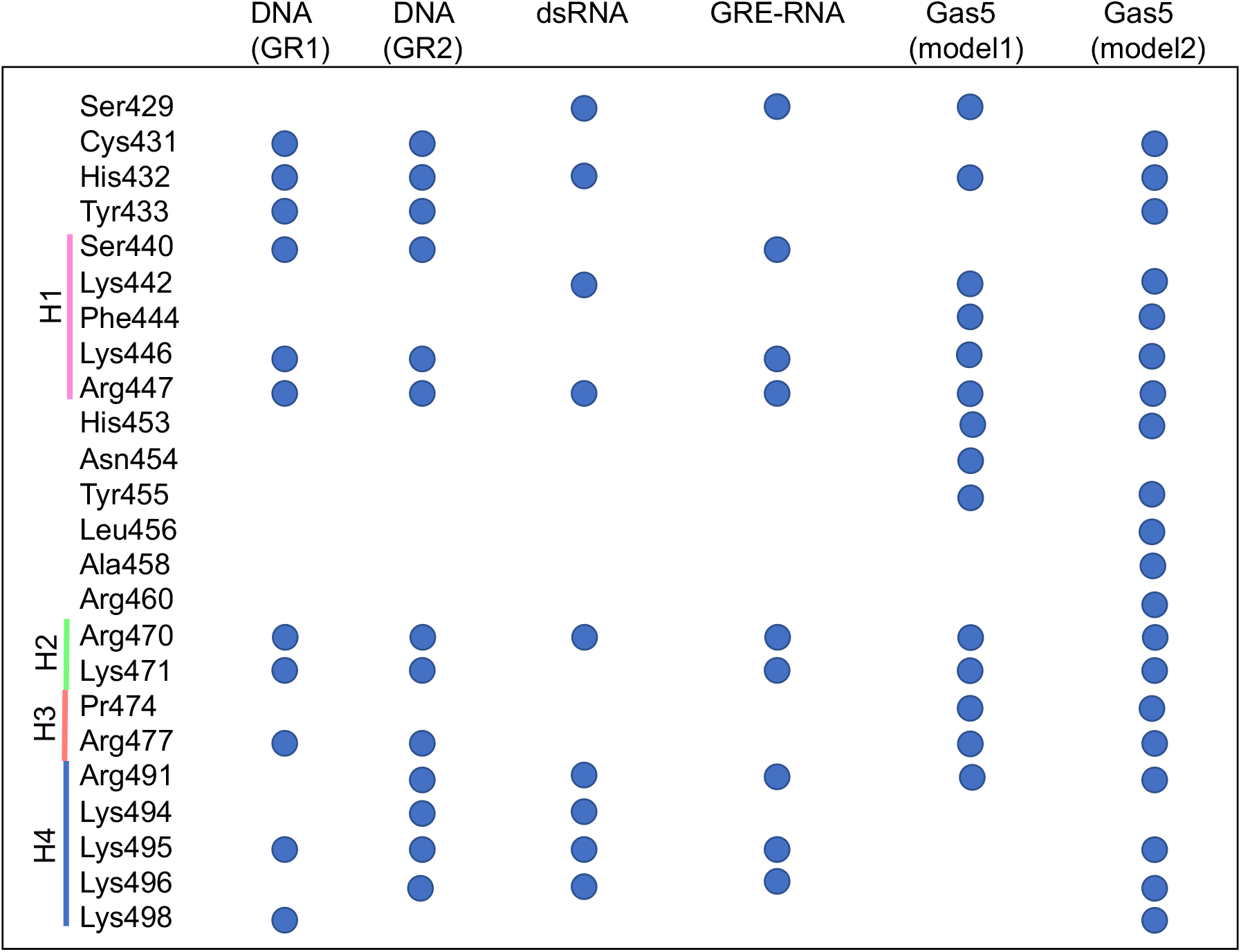
Residues of the glucocorticoid receptor involved in nonspecific interactions with different nucleic acids model systems.

In case of the GR–RNA binding, our dynamic contacts maps show, in agreement with the experiment (4), the dominating role of the electrostatic nonspecific interactions for the GR-RNA complex formation (Figures 3B-E, 4). Though, depending on the RNA structure, GR may exhibit some specific contacts as well (Figures 3B-E, 4). We would like to clarify that by the “specific” interactions we here mean the contacts formed between the RNA bases and the GR sidechains, exchanging a “specifically” interacting RNA base to another would have no or little impact on the binding affinity. Specific GR-RNA contacts involve the protein residues of helix 4 and the flexible nucleotides within the terminal loops of the GRE_RNA and Gas5_RNA hairpins. Additional specific contacts include, for GR-GRE_RNA, the contact by Lys442 of the recognition helix to C16w of the terminal loop, for GR-Gas5_RNA – by His453, Asn454 (Gas5_model1) and Leu456 (Gas5_model2) of the region between the recognition helix and helix 2 to the U28-A29 bulge bases. The latter specific GR-Gas5-RNA contacts become possible due to the presence of the bulge, which allows the RNA molecule to bend toward the protein (see Groove and Helical Parameters for further details).

Overall, we observe that all the residues of the recognition helix, except Val443 (and Lys442 for GRE_RNA) that interact specifically with DNA, form nonspecific contacts with RNA. Also, most of the residues involved in the nonspecific contacts with RNA, exhibit nonspecific interactions with DNA (Figures 3-4). The only unique nonspecific interactions, seen for the GR-Gas5_RNA system, include the GR residues 453-458 interacting with the Gas5_RNA bulge. Therefore, based on the analysis of the GR-nucleic acids contacts, we do not observe any specific set of GR residues, uniquely involved in the RNA binding. We would like to clarify that our observation is limited to the GR-DBD residues present in the employed crystal structure (PDB ID, 6CFN, a. a. r. 418-500).

The alanine scanning experiments (4) have revealed a significant role of helix 4 for the GR-RNA associations with the following contribution order: Lys492>Arg491>Lys495>Lys494> Lys496>Lys498. Our simulations show a preference for the Arg491-RNA interactions over the Lys494-RNA interactions. Lys492 is observed to electrostatically interact with GRE_RNA in less than 4% of the corresponding MD trajectory snapshots. Our simulations also indicate that it is important for helix 4 to unfold into a random-coil structure to be able to form more electrostatic interactions with RNA. For the GR-GAS5_RNA_model1 system, helix 4 does not unfold due to the nucleotide fraying within the terminal loop; this results in a fewer helix 4-Gas5_RNA interactions. Here we would like to point out that sampling of all conformational substates exploited by flexible random-coil regions require significantly longer simulations, beyond the μs range, which could explain why we do not see long-lived Lys492-RNA interactions in our simulations. The role of Lys492 could also be to impact the flexibility of helix 4, instead of directly interacting with RNA.

The results from our simulations point toward an importance of both the recognition helix and helix 4 for the GR-RNA interactions, where the residues within these two regions allow GR to anchor itself within the RNA molecule. Experiments show that GR exhibit poor binding affinity towards dsRNA (Kd>3500 nM). Our simulations show that nonspecific interactions exploited by the recognition helix (Lys442 and Arg470) but also residues outside this region (Ser429, His432 and Arg470) are lost toward the end of the GR-dsRNA MD trajectory (Figure S6). The recognition helix cannot follow the changes of dsRNA major groove geometry (see Groove and Helical Parameters for further details). The only few GR-dsRNA contacts left towards the end of the trajectory are those exploited by unfolded helix 4(Figure S6).

### Protein–Nucleic Acids Interaction Energies and Configurational Entropies

The binding experiments (4) show the following GR binding preference order: DNA > Gas5_RNA > GRE_RNA >>> dsRNA. To explore if our simulations capture the same trend, we next analyse the intermolecular interaction energies (Figures 5, S9-10). Here we would like to point out that the accurate calculation of protein-DNA/RNA interaction energies is challenging due to the massive negatively charged backbone of nucleic acids. Nevertheless, to see if we can capture a qualitative trend, we calculate the interaction energies including electrostatic and van der Waals components, for the specific and nonspecific protein-DNA/RNA contacts along all protein-bound trajectories with GROMACS energy analysis tool. The GR-DNA complex shows more favourable interaction energies for the specific contacts, whereas for the nonspecific contacts the energies are similar to those for the GR-RNA complexes. The cumulative interaction energies, going from more negative to less negative energies, show the following trend: GR-Gas5_RNA_model2 >> GR(mon1)-DNA > GR(mon1)-DNA > GR-Gas5_RNA_model1 ≈ GR-GRE_RNA >> GR-dsRNA. The trend is consistent with the one observed experimentally. Our simulations further detail that the unfolding of helix 4 allows for the most favourable GR-Gas5_RNA interaction energies, if compared to the interaction energies of GR-DNA monomeric complexes. However, the GR binding to DNA is dimeric. Thus, combining the protein-DNA interactions energies for both GR monomers, shows a stronger GR preference for DNA over RNA.

**Figure 5.**
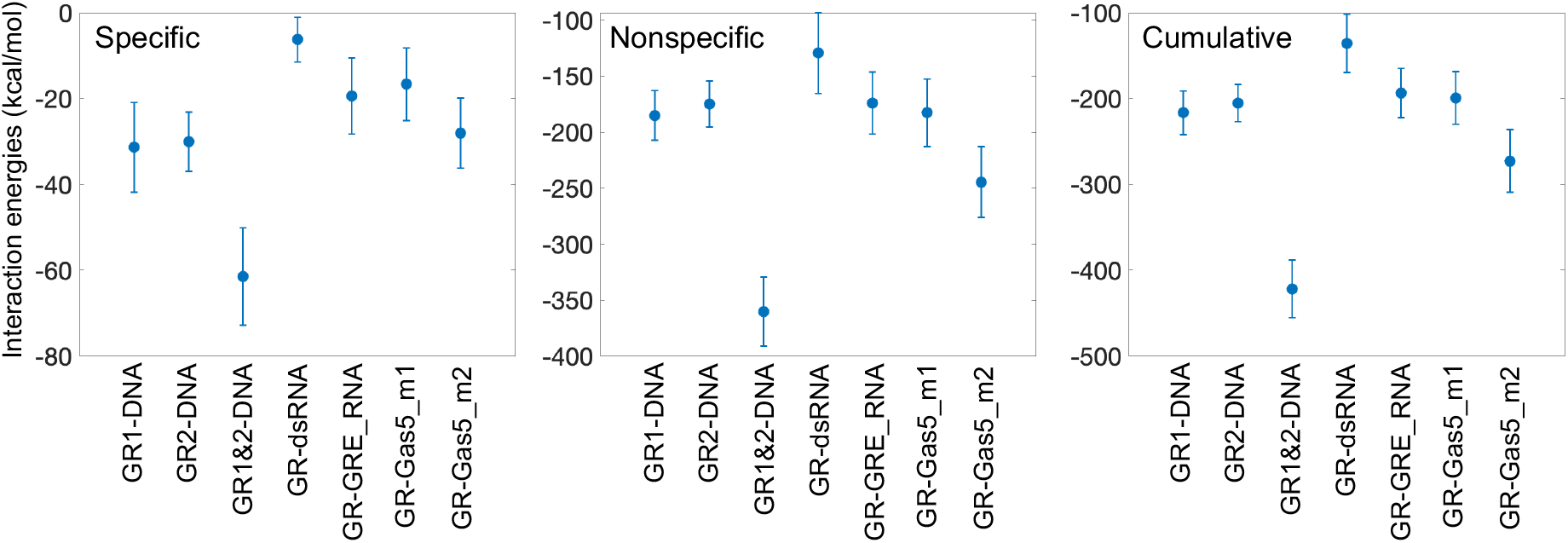
Protein-DNA/RNA interaction energies (kcal/mol), calculated for all GR-DNA/RNA complexes.

We also calculate the configurational entropies for all GR-nucleic acids systems to explore further the thermodynamics of the binding process. The configurational entropy is a part of the total entropy change, which arises from the degrees of freedom of the solute. NMR experiments have shown that for proteins the configurational entropy and the solvent entropy contributions can be similar in magnitude (54, 55), and thus can impact the thermodynamics of protein interactions. We calculate the configurational entropies using the Schlitter’s formula (56) in the presence and absence of GR for the entire DNA/RNA sequence and only for the response element (RE). To estimate the standard deviations, we calculate the configurational entropies for the entire 500 ns trajectory and for all consecutive 100 ns windows within. Upon the GR binding to DNA/RNA, we expect the configurational entropy to decrease if the two molecules are structurally complementary and form a stable complex. As if the protein makes the nucleic acid molecule more rigid for global motions. Looking at the configurational entropies for the entire sequences (Table 1), we see that this statement is true for GR-bound DNA and both Gas5 systems, where the configurational entropies decrease by ∼15, 5 and 7.5 kcal/mol, respectively. In the case of GRE_RNA, the decrease in insignificant, 0.37 kcal/mol. While, for GR-bound dsRNA the configurational entropy remains approximately the same (increased by 0.014 kcal/mol). Similar situation occurs through the analysis of the configurational entropies of the RE-sites, where the decrease in the configurational entropy upon the GR binding to dsRNA is ∼10-100 times smaller than in other systems. These results further support that the GR-dsRNA complex is less stable, and the protein cannot adjust to the binding site on dsRNA.

**Table 1:** Configurational entropies [TS (kcal/mol), for T=300K] for the DNA/RNA backbone P atoms

|  | Whole sequence<br>(kcal/mol) | Response Element<br>(kcal/mol) |  |
| --- | --- | --- | --- |
| DNA | 67.41±1.37 | 13.48±0.364 (RE1) | 12.65±0.370 (RE2) |
| GR-DNA | 52.66±1.83<br>(-14.75) | 10.03±0.521 (RE1)<br>(-3.45) | 10.07±0.771 (RE2)<br>(-2.58) |
| dsRNA | 51.98±0.502 | 9.604±0.0551 |  |
| GR-dsRNA | 52.00±0.560<br>(0.0138) | 9.573±0.109<br>(-0.0306) |  |
| GRE_RNA | 37.45±0.234 | 9.999±0.0527 |  |
| GR-GRE_RNA | 37.08±0.775<br>(-0.367) | 9.362±0.394<br>(-0.637) |  |
| Gas5_model1 | 46.38±0.774 | 9.813±0.116 |  |
| GR-Gas5_model1 | 41.41±1.36<br>(-4.96) | 9.440±0.228<br>(-0.372) |  |
| Gas5_model2 | 47.16±2.36 | 10.16±0.314 |  |
| GR-Gas5_model2 | 39.63±1.73<br>(-7.54) | 8.656±0.299<br>(-1.50) |  |

*Note: Entropies have been derived for the windows: last 500ns, 100-200ns, 200-300ns, 300-400ns and 400-500ns to obtain the standard deviations. Values in parathesis are the differences in mean values with respect to the naked nucleic acid molecule. RE1 and RE2 are the two response elements presence on the DNA molecule.*

### Nucleic Acids Helical and Groove Parameters

Next, we analyse changes in helical and groove parameters upon the GR binding for all studied nucleic acids systems (Figures 6-7 and Figures S11-18). We have previously shown that local changes in DNA groove geometries and helical parameters, mainly in shift and slide, facilitate the direct-read-out mechanism (49, 57); that is the ability of proteins to form specific contacts with their genetic sites. The binding of GR to DNA, as expected, brings in alterations in shift, slide and twist within the GRE-sequence and the flanking sites (Figures 6, S11). Additionally, the GR binding also leads to small changes in roll and tilt angles (Figures 6, S11), due to the hydrophobic interactions of the bulky Val443 residue and the interactions of helix 4 with DNA minor groove. Furthermore, the GR monomers make DNA major groove wider within the GRE-sites, and narrower within the linking region between the two monomers (Figures 6, S15). Several b.ps. within the GRE-site shift towards the major groove, as illustrated by changes in x-displacement (Figure 6), making the major groove shallower and more accessible for specific interactions with the protein.

**Figure 6.**
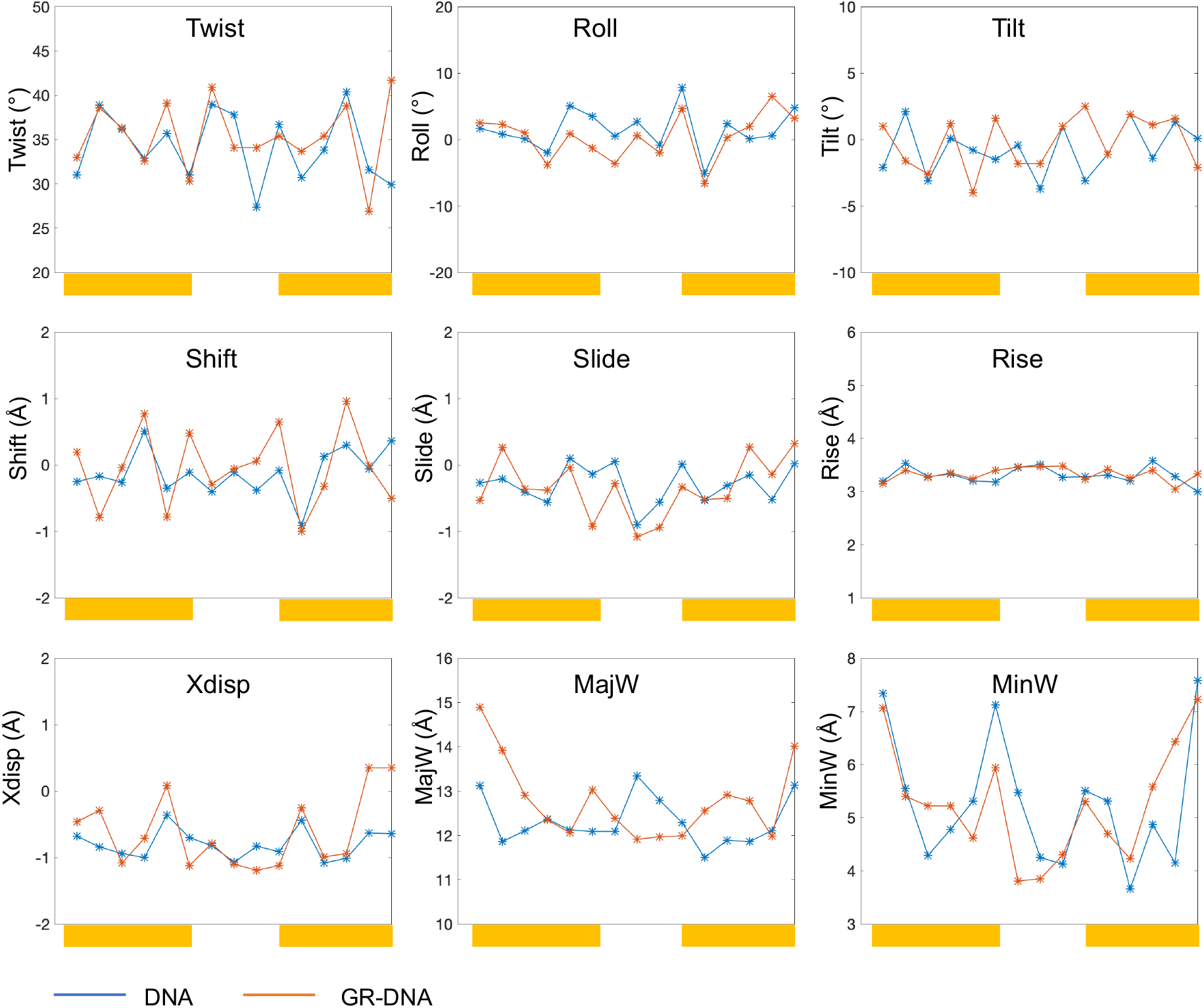
Average helical and groove parameters for unbound (blue) and GR-bound (orange) DNA. The binding sites for the two GR monomers are highlighted with yellow rectangular boxes.

**Figure 7.**
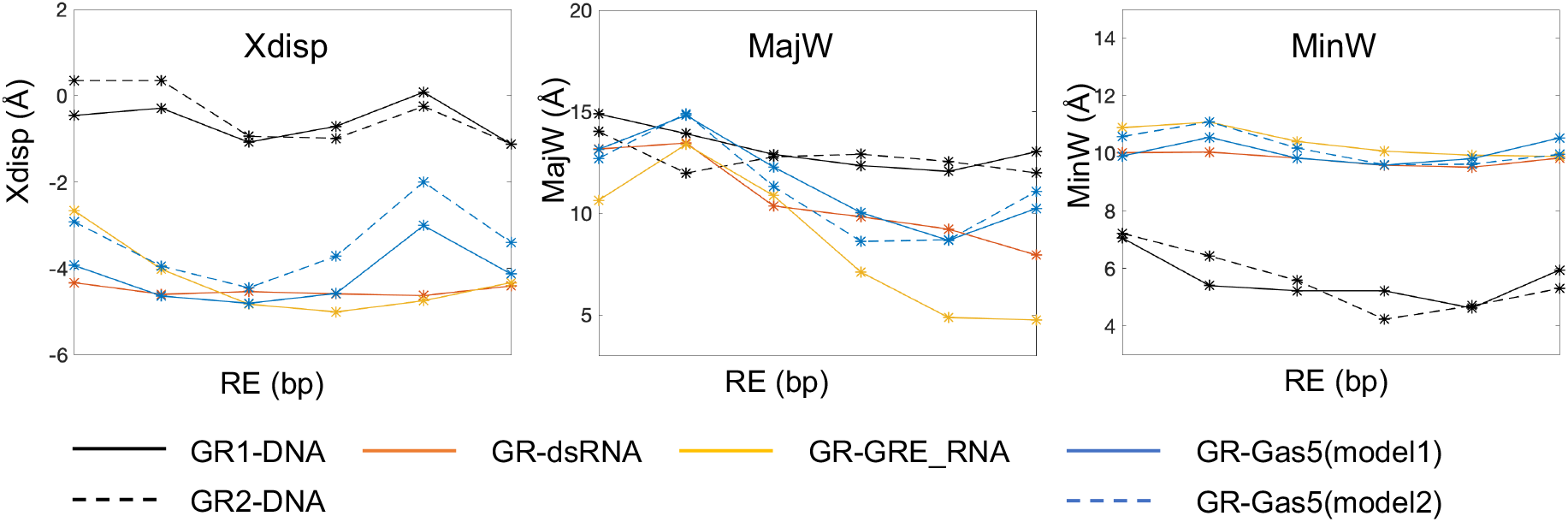
Comparison of average values for x-displacement and groove widths in different nucleic acids model systems for the b.p. within the GRE-site for the GR-bound systems.

In contrast to DNA, the binding of GR to the RNA systems induces no notable changes in helical parameters (Figures S12-S14). This is not surprising as the helical parameters distributions appear monomodal within the GRE-site for all unbound RNA, which means that the RNA b.ps. are quite rigid and the GR residues cannot reach them to form specific contacts. Also, the values of helical parameters for the GR-bound RNA molecules (except rise) are distinct from those of GR-bound DNA (Figure S19). Nevertheless, we observe that the presence of a hairpin and a bulge result in different values of helical parameters for GR-bound RNA, within the GRE-site (Figure 6). Most notably, for GR-bound Gas5_RNA the last 3 b.ps. of the GRE-site (AGA**CUG**), adjacent to the bulge, have the least difference in twist, shift, tilt, and x-displacement values (Figure 7) with respect to the same parameters of DNA. For GR-bound GRE_RNA, the only significant difference from dsRNA is in the higher values of b.ps. roll angle.

The GR binding also has little impact on the RNA systems grooves geometries. The RNA grooves geometries differ depending on the structural motifs of the RNA systems (Figures S16-18). When comparing the major groove width within the GRE-site across all studied GR-bound nucleic acids systems, we notice that the first 3 b.ps. exhibit comparable values (Figure 6), which suggest a suitable binding pocket for the protein initial binding. For GRE_RNA and Gas5_RNA, by the first 3 b.ps. we refer to the ones on the hairpin side. However, it should be noted that the major groove width distributions for dsRNA are broader (Figures S16-S18), which indicates larger structural fluctuations, which contribute to the instability of the GR-dsRNA contacts network and thus low binding affinity. The major groove widths of the remaining 3 b.ps of the GRE-site differ in the RNA systems, with Gas5_RNA being the closest to DNA, where we refer to the Gas5_RNA GRE-half site adjacent to the bulge. b.ps. The conformational similarities between Gas5_RNA and DNA, we believe, contribute to the more favourable protein-Gas5_RNA contacts network, interaction energies and configurational entropy changes among the studied RNA systems.

### Principal Component Analysis

We perform principal component analysis (PCA)(58, 59) of the trajectories to see if there are any predominant molecular motions that could explain the GR preference for DNA and hairpin RNA over dsRNA. First, we do the analysis for the unbound nucleic acids systems. The most dominant motions in the MD trajectories are represented within the first three principal components, which accounts for ∼60-65% of the total variance (Table S1, Figure S20A).

For DNA, dsRNA, and GRE_RNA the first three components describe similar motions, where the first two components correspond to sideward and upward bending, and the third component describes the twisting rotation (Figure S21A-C, Movies S1-9). The three motions appear to impact the open state of the major groove. However, the structural dynamics of the GRE-site binding pocket differs in the three systems. In DNA the shape fluctuations of the GRE-site binding pocket are small, whereas in dsRNA the pocket becomes extremely deep and narrow. In GRE_RNA the binding pocket appears smaller and tilted to one side of the major groove, explaining the observed asymmetry of the GR binding.

For Gas5_RNA systems the motion described by the first component dominates. It involves a bending motion that changes the size of the major groove pocket, going from an open to a closed and bent conformation (Figure S21D-E, Movies S10 and S13), where the middle state resembles the averaged structure of the GR-bound state. The motions described by the second and third components differ between the Gas5_RNA_model1 and model2. In model1, they correspond to a rotation and a downwards bending motions. Contrary, in model2 the second and third components describe a rotational bending motion where several terminal loop residues rearrange to make the binding pocket more open (Figure S21D-E, Movies S11-12, S14-15).

We continue with the PCA analysis of the trajectories for the protein-bound systems (Figures S20B-D, S22-23). We do the analysis in three ways; (1) for the heavy atoms of the protein-DNA/RNA systems and (2) for heavy atoms of DNA/RNA with the superposition on the nucleic acids in both cases; and (3) for the heavy atoms of the complexes with the superposition on the entire protein-DNA/RNA systems. The three analyses are complementary: (1) highlights the GR movements upon changes in the RNA/DNA structures; (2) – conformational changes in RNA/DNA upon the protein binding; (3) – combined conformational changes for the protein-DNA/RNA complexes. It must be noted that caution should be taken when interpreting the results of analysis (3) as finding the best least-square fit for the entire complex is nontrivial and the analysis may provide artefacts. Analysis (3) for all studied GR-RNA/DNA systems reveals no significant global conformational changes (Movies S22-24, S31-33, S40-42, S49-51, S58-60), and thus will be omitted from further discussions.

For all protein-bound systems, the first three principal components from the three analyses account for ∼50-70% of the total variance (Table S1, Figure S20B). For the GR-DNA system, analysis (1) reveals that both GR monomers are stably bound to DNA. The GR dimer makes DNA more rigid, and the three principal components correspond to small sideward bending motions (Figures S22A, Movies S16-18), which the GR dimer follows. The largest amplitude movements are seen for helix 4 of GR, which correspond to swinging out of DNA minor groove. Analysis (2) shows motions described by the first and third components similar to those of unbound DNA, whereas the second component shows an expansion of DNA major groove as a result of the GR dimer binding (Figure S23A, Movie S19-21).

For GR-dsRNA system (Figure S22B, Movies S25-27), analysis (1) reveals a translational movement of the protein: the first component describes a sidewards bending of dsRNA, which closes the major groove and makes GR tilt and shift from one side of the GRE-site to the other. The second component involves the opening of the major groove, which allows GR to sink into to the GRE-pocket, but not as deeply as is in the GR-DNA complex. The third component describes a transition to a closed major groove state through bending and the loss of contacts of the GR recognition helix with dsRNA. Analysis (2) (Figure S23B, Movies S28-30) results in the first two components describing a downward and sideward bending that impact the open state of the major groove, which forces GR to shift to remain bound to dsRNA. The third component corresponds to the movement, similar to that of the first two components of analysis (1). Comparing the motions from PCA analyses (1) and (2), and the ones of the unbound state, we conclude that the protein stimulates the opening of the dsRNA major groove. However, the dsRNA molecule cannot maintain a suitable width of the binding pocket for the GR monomer, resulting in the protein dissociation.

For the GR-GRE_RNA and both GR-Gas5_RNA systems (Figures S22-23C-E, Movies S34-39, S43-48, S52-57), similarly to GR-DNA, analysis (1) shows the protein stably bound to RNA. However, we observe different predominant motions in the three RNA hairpin systems, and as a result a different structural response of GR. For GRE_RNA, we observe breathing upward and sideward bending motions that impact the compactness of the GRE-site, leading to the rocking and tilting motions of the recognition helix and helix 4. For Gas5_RNA_model1, the motions described by the first three principal components resemble those for the GRE_RNA case, but the amplitude is bigger suggesting a more open and flexible structure. The GR protein adjusts by tilting and turning of its recognition helix to follow the changes in the shape of the binding pocket. The motions of the Gas5_RNA_model2 also include sidewards bending, but in this case, additionally we observe the expansion of the minor groove adjacent to the GRE-site. This motion allows residues from GR helix 2 to approach the groove and interact with the bulge region, leading to a forward tilting movement of the GR recognition helix. Uniquely, for Gas5_RNA_model2, the sideward bending, captured by the component 3 of analysis (1) (Figures S22-23, Movie S54), also involves movements of the terminal loop and bulge residues.

Overall, the PCA analysis confirms that to anchor within the GRE-binding site, the protein requires a wide and shallow major groove pocket. The protein effectively moulds the GRE-site within DNA major groove but has little effect on the major groove geometry of the RNA systems. Furthermore, to remain bound the protein needs to adjust to the structural dynamics of RNA through changes in the orientation of the recognition helix, which in turn requires a more open conformation of the GRE-site supported by the terminal hairpin. Addition of the bulge, in Gas5_RNA, increases the kinking motions of RNA and provides an additional interaction surface for GR helix 2.

## CONCLUSIONS

Using all-atomistic MD simulations, we explore the mechanistic details of transcription factor associations with RNA. As the model system we employ the glucocorticoid receptor (GR) bound to several nucleic acids model systems, which include the native DNA target and three structurally different RNA molecules containing the glucocorticoid response element (GRE) and GRE-like sequences. The studied RNA systems include dsRNA, a fully paired RNA hairpin, and an RNA hairpin with a bulge. The latter system is derived from long-noncoding Gas5 RNA, which was shown to interact with GR *in vivo* (16, 18). Through the analyses of the protein-DNA/RNA contacts networks, interaction energies, configurational entropies, nucleic acids helical and groove parameters, and principal component analysis, we conclude that GR DNA-binding domain (DBD) can bind both DNA and RNA, but the association mechanisms are different.

The DNA recognition by a GR dimer involves formation of several specific contacts between the residues of the GR-DBD recognition helixes and the bases of the GRE-sites. These specific contacts anchor each recognition helix deep within DNA major groove, leading to the significant widening of the major groove of both GRE-sites (Movie S20). It is exactly one DNA turn that separates the centers of two GRE-sites, which allows for the strongest allosteric communication between the monomers of the GR dimer (60, 61), explaining the observed positive cooperativity for the GR-DNA association (4, 28).

Contrary, the GR-DBD-RNA complexation depends predominantly on nonspecific electrostatic contacts. Though, the number and strength of these nonspecific contacts depend on the structural motif of the RNA molecule (Figures 3, S3-S6). A GR-complex with uniform dsRNA, with its characteristic A-form double helix, shows fewer contacts, mostly from the residues of helix 4. The GR-dsRNA MD trajectory shows a gradual loss of protein-RNA contacts and the beginning of the protein dissociation. Contrary, a hairpin motif allows RNA to adapt a more open conformation with a rather wide and shallow major groove pocket, allowing for more GR-RNA contacts to be formed. The GR interacts with the Gas5_RNA and GRE_RNA hairpins through contacts formed by the positively charged residues of the recognition helix and helix 4. Addition of the internal loop (bulge) of Gas5_RNA increases flexibility of the off-GRE stem, which contributes with additional contacts formed by the residues adjacent to the GR dimerization motif.

The binding energies and the configurational entropies analyses of the GR-DNA/RNA complexes show the same binding affinities relations, as the binding experiments (4): DNA > Gas5_RNA > GRE_RNA >>> dsRNA. In addition, computationally we can estimate the binding energies for each GR monomer in the GR-DNA complex, which further reveals that a GR monomer can bind Gas5_RNA even stronger than DNA. This is possible due to the combination of two factors. The first being the above-mentioned GR-Gas5_RNA nonspecific interactions, because of the increased flexibility of the off-GRE stem. The second being the unfolding of helix 4, the residues of which then interact specifically with the bases of the terminal loop region. Crystal structures support that helix 4 can exist as both folded (36, 62) and unfolded (28, 63). In our computational experiments, we observe the unfolding of helix 4 for GR-DNA, GR-GRE_RNA, and GR-Gas5_RNA_model1 systems. Though, we believe that the unfolding of helix 4 in the GR-DNA MD trajectory is the consequence of GRE-flanking regions being too short. The unfolding of helix 4 may present a mechanism for the GR-nucleic acids structural recognition – a hypothesis that we plan to investigate further in the future.

Taken together, we demonstrate with atomic level detail that the glucocorticoid receptor employs the sequence recognition mechanism when binding DNA and the shape recognition mechanism when binding RNA. The protein interacts with either DNA or RNA using nearly same residues, yet the nature of the formed protein-nucleic acids contacts and the binding interfaces differ. We believe that the described dual recognition mechanism may be shared by other zinc finger transcription factors.

## Acknowledgements

The authors would like to dedicate this work to late Dr. Matti Lepistö, who laid the foundation for this project. The authors thank Prof. Deborah S. Wuttke and Prof. Robert T. Batey for insightful discussions, and Swedish National Infrastructure for Computing (SNIC) for the generous provision of computing resources.

## Author Contributions

JH and AR conceived and designed the study. JH performed the MD simulations. JH and AR analysed the data. JH and AR wrote the article.

## Financial Support

Swedish Foundation for Strategic Research SSF Grant [ITM170431] and Magn. Bergvalls Foundation Grant to A.R.

## Conflicts of Interest declarations

### Conflicts of Interest

None

## Data availability statement

Generated datasets from analyses of MD trajectories, including protein–nucleic acids contacts information, interaction energies, helical parameters of unbound and protein–bound nucleic acids; also starting, last snapshot, and average structures for unbound and protein–bound nucleic acids are freely accessible at zenodo.org as https://doi.org/10.5281/zenodo.7738133.

## Notes

https://doi.org/10.5281/zenodo.7738133

### Competing Interest Statement

The authors have declared no competing interest.

