## Supplementary information for "Decoding the Dual Recognition Mechanism of Glucocorticoid Receptor for DNA and RNA: Sequence vs. Shape"

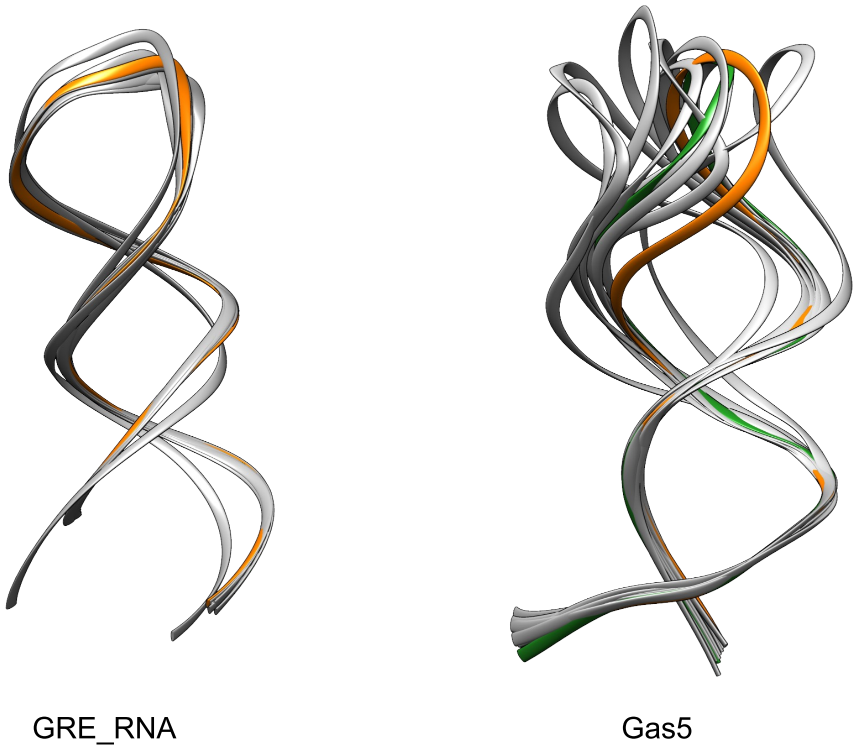


**Figure S1:** Derived models of GRE_RNA and Gas5_RNA. Models highlighted with orange (the highest ranked model) and green (the second highest ranked model) were used for the docking studies.

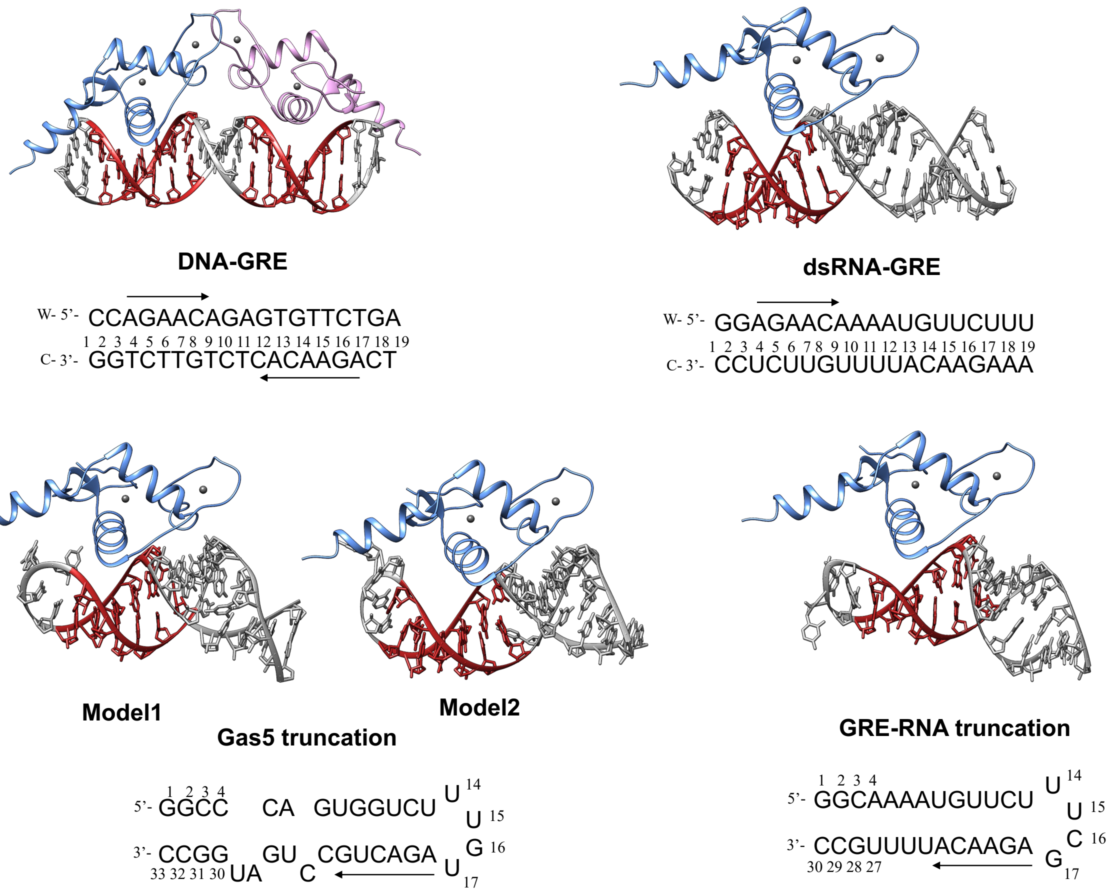


**Figure S2:** Prepared GR-DNA/RNA complexes.

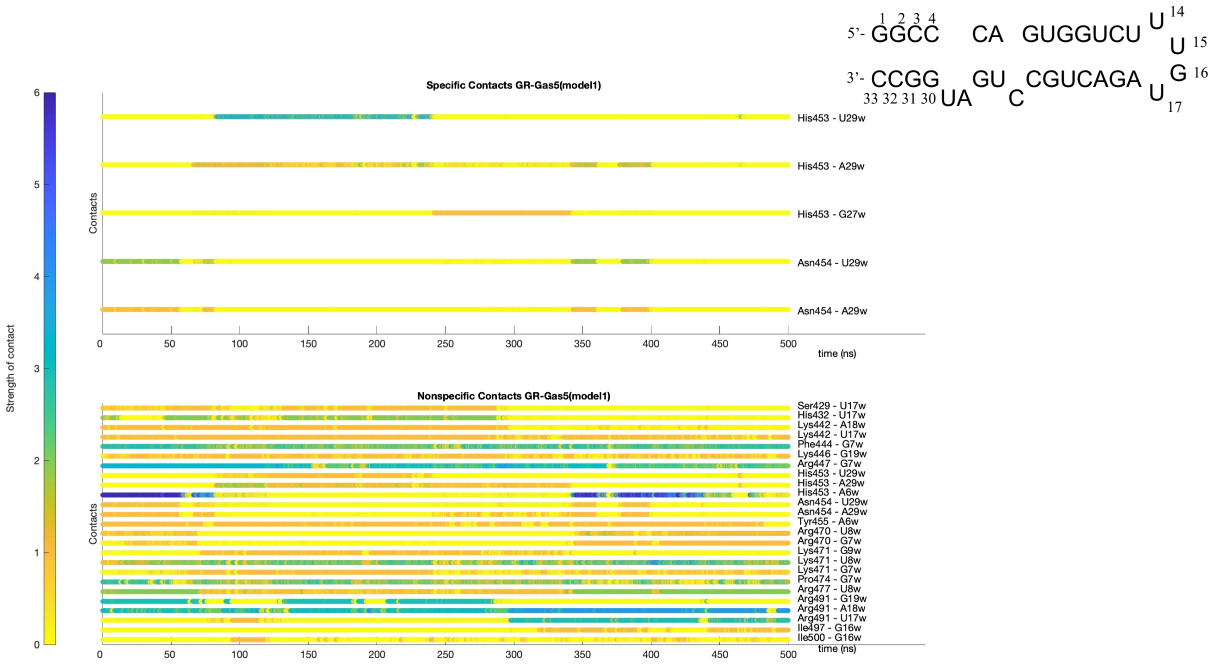


**Figure S3:** Dynamic contacts map for the specific and nonspecific GR-Gas5_RNA_model1 interactions.

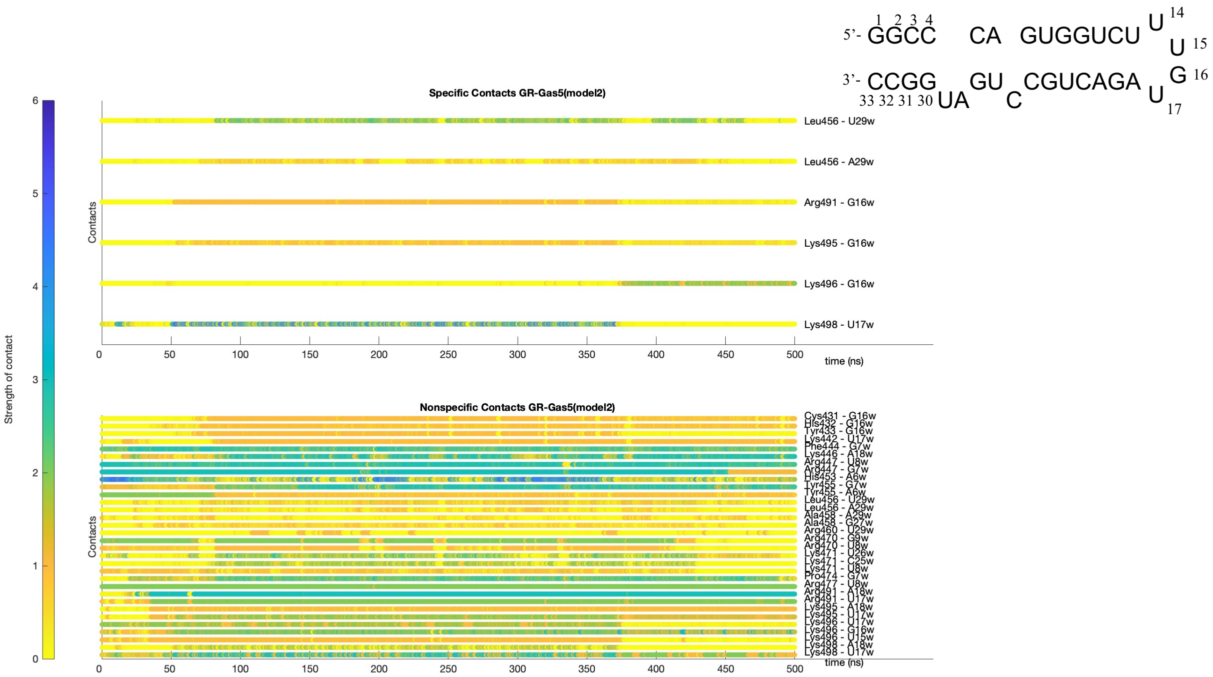


**Figure S4:** Dynamic contacts map for the specific and nonspecific GR-Gas5_RNA_model2 interactions.

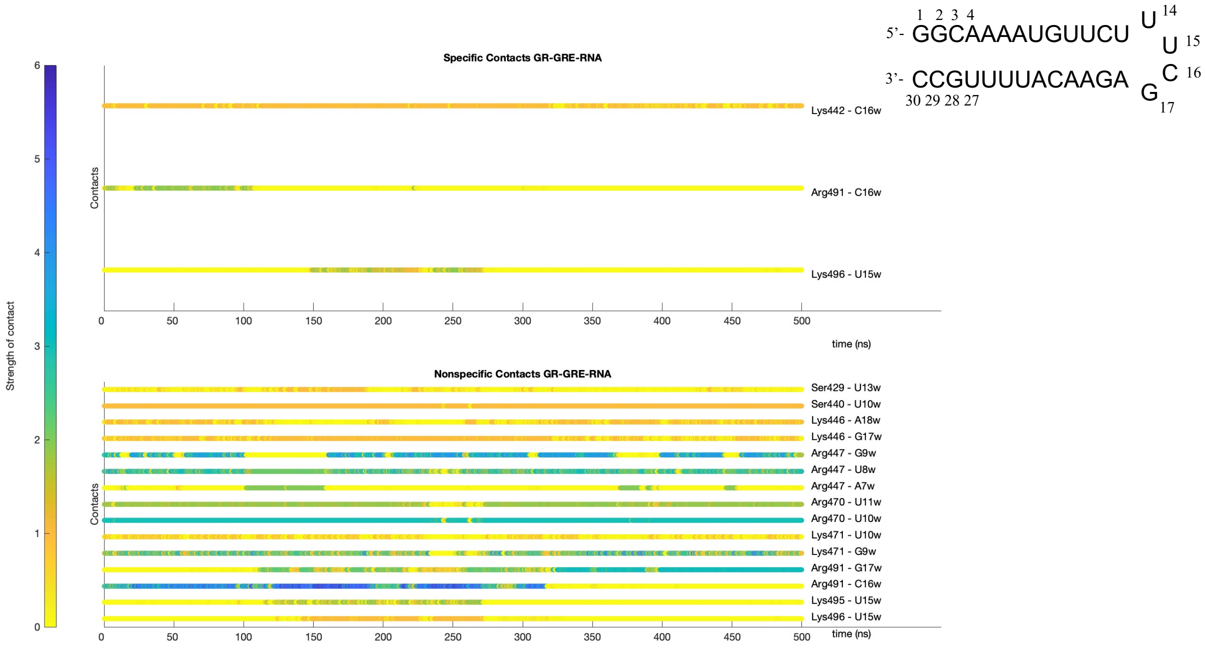


**Figure S5:** Dynamic contacts map for the specific and nonspecific GR-GRE_RNA interactions.

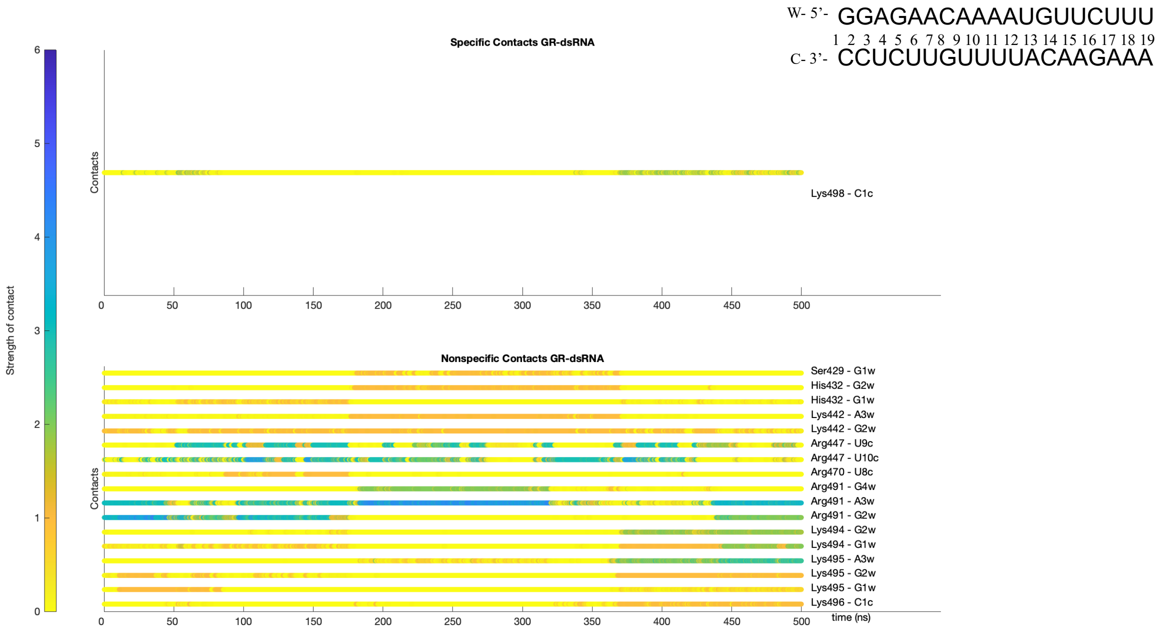


Figure S6: Dynamic contacts map for the specific and nonspecific GR-dsRNA interactions.

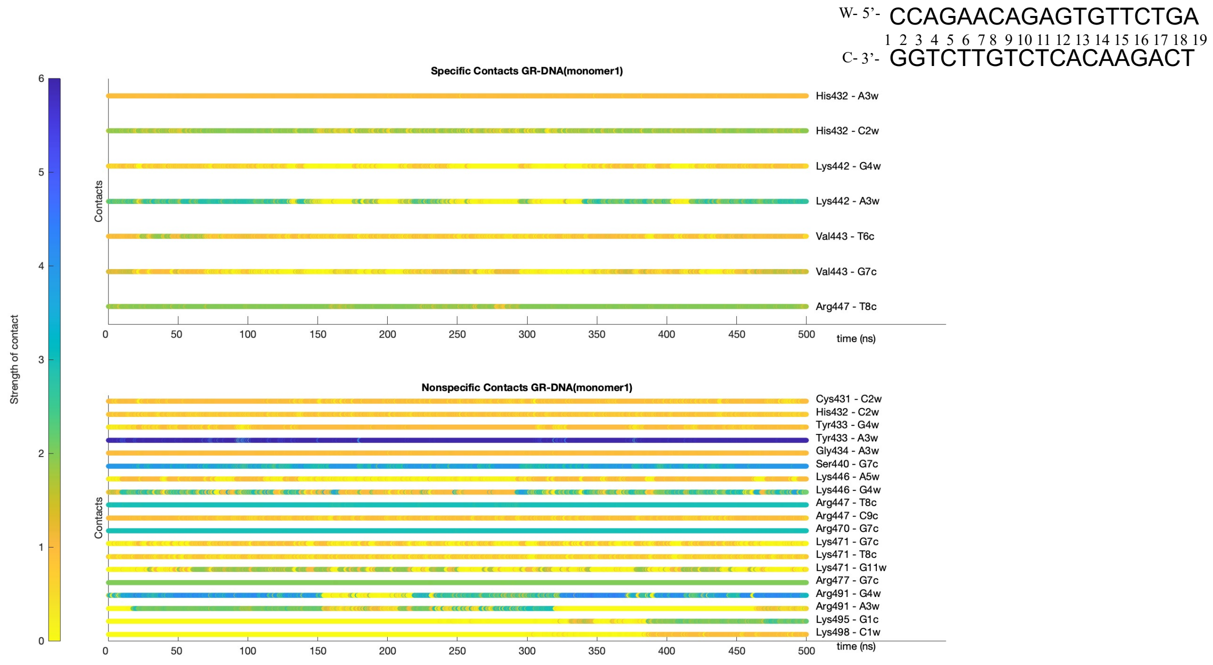


**Figure S7**: Dynamic contacts map for the specific and nonspecific GR(monomer1)-DNA interactions.

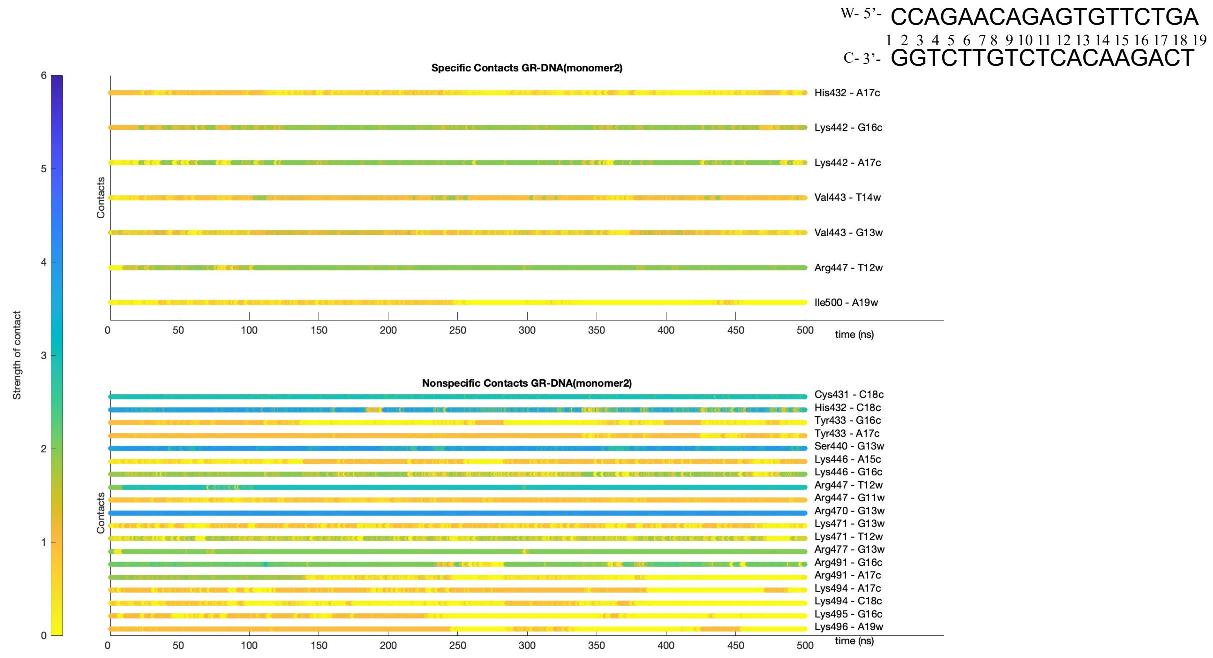


**Figure S8:** Dynamic contact map for the specific and nonspecific GR(monomer2)-DNA interactions.

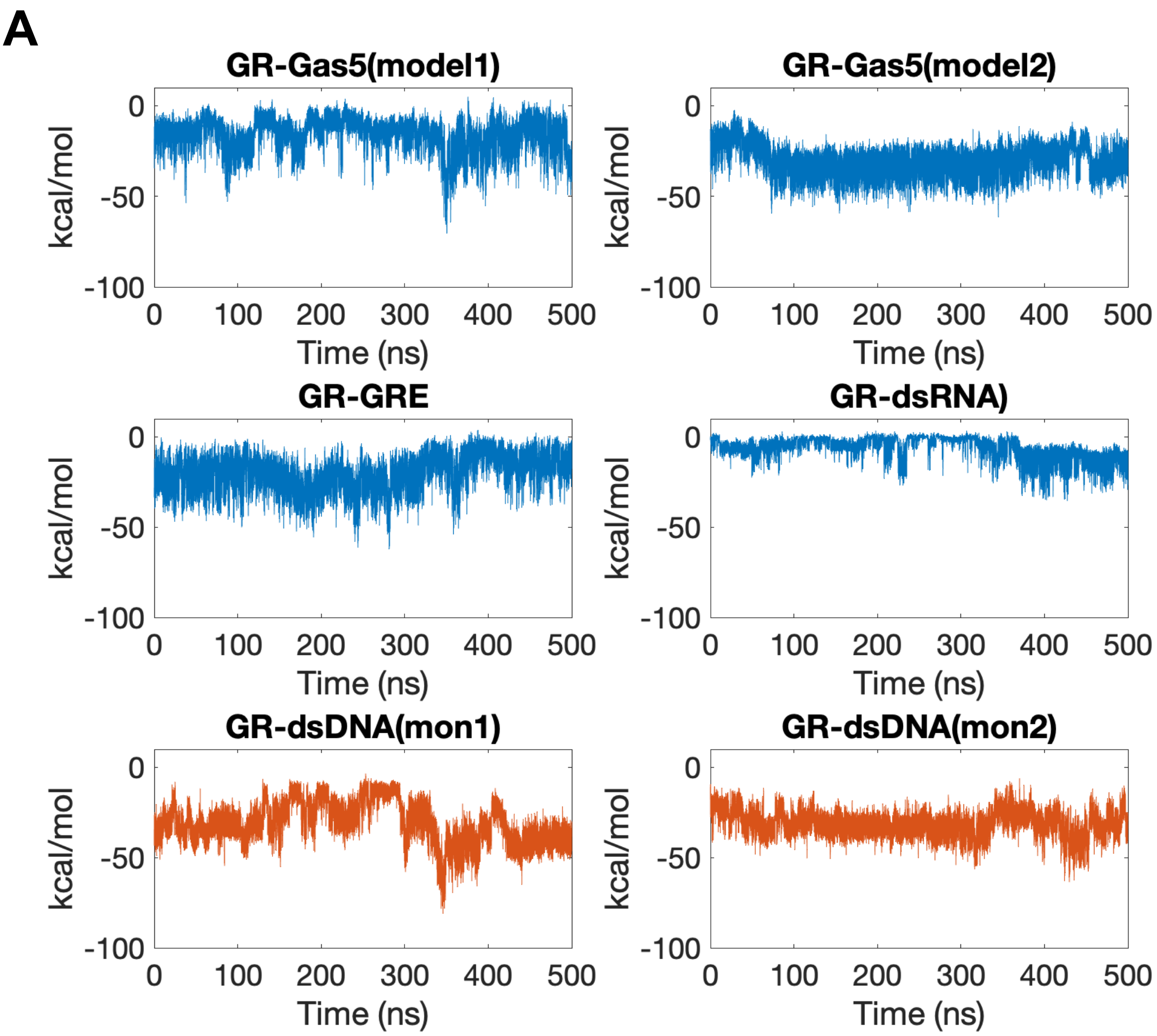
A
GR-Gas5(model1)
GR-Gas5(model2)
GR-GRE
GR-dsRNA)
GR-dsDNA(mon1)
GR-dsDNA(mon2)
kcal/mol
0
-50
-100
0
100
200
300
400
500
Time (ns)

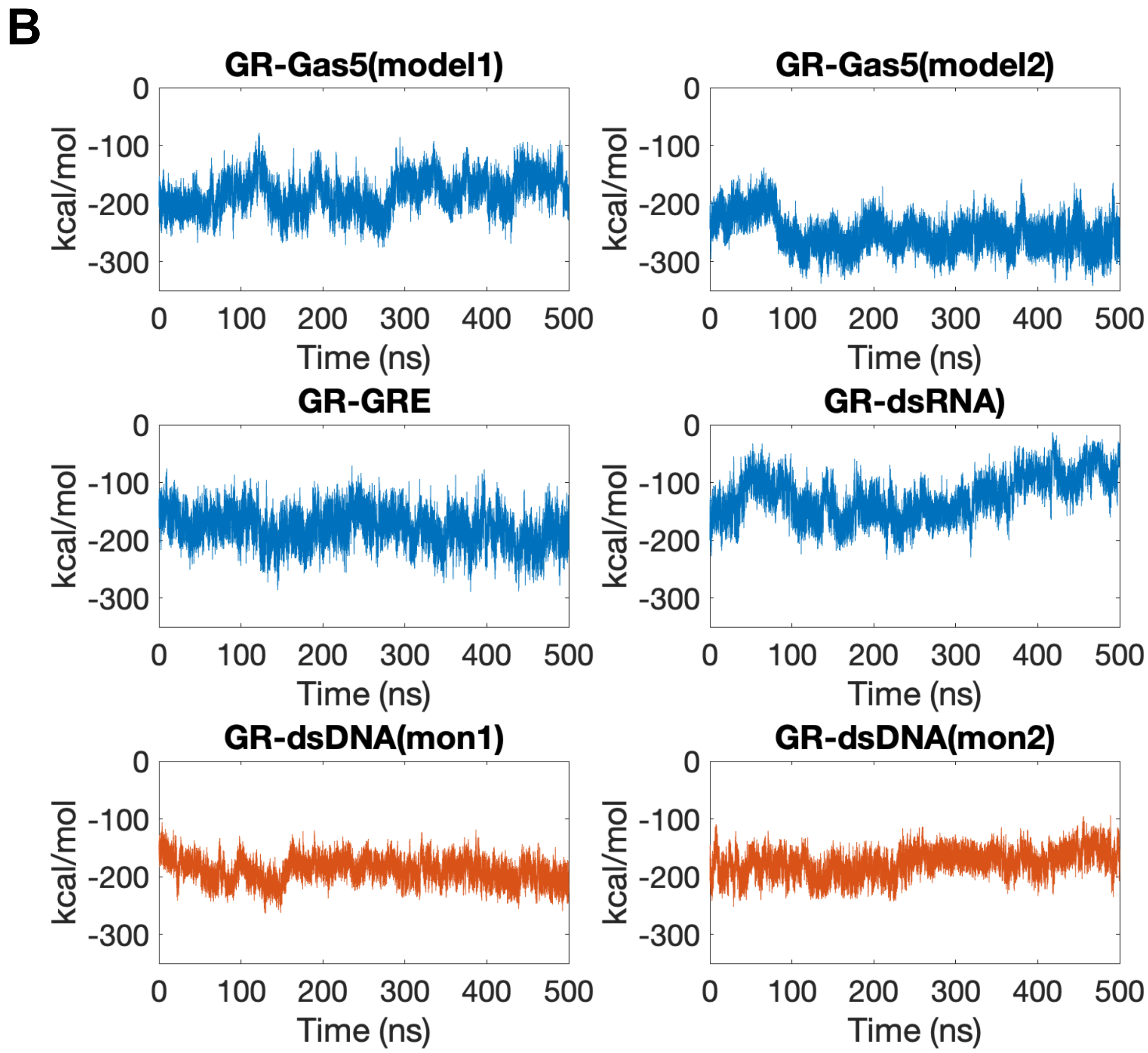


**Figure S9:** Specific (A) and nonspecific (B) GR-DNA/RNA interaction energies along the corresponding MD trajectories.

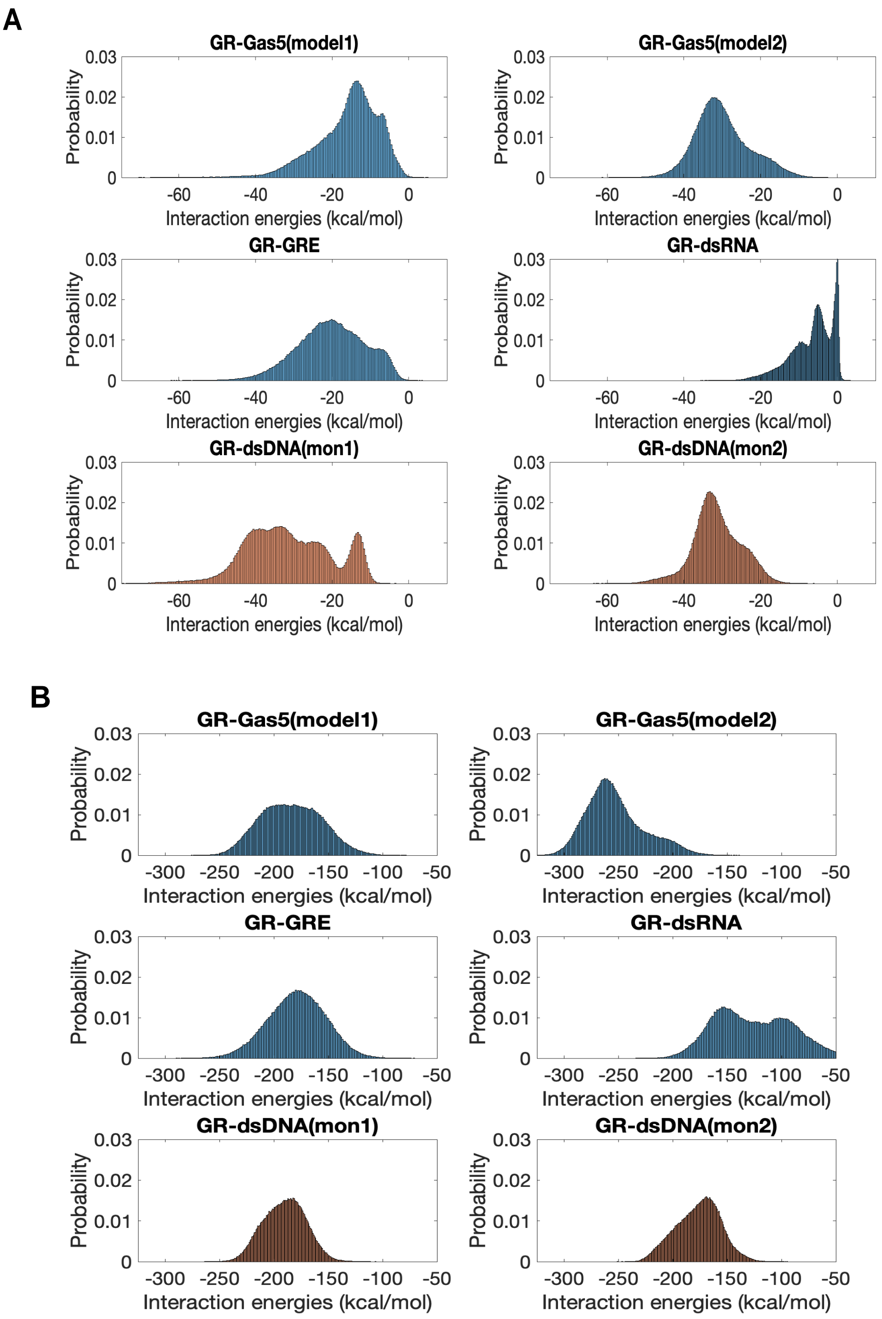


**Figure S10:** Specific (A) and nonspecific (B) GR-DNA/RNA interaction energies distributions

**A**

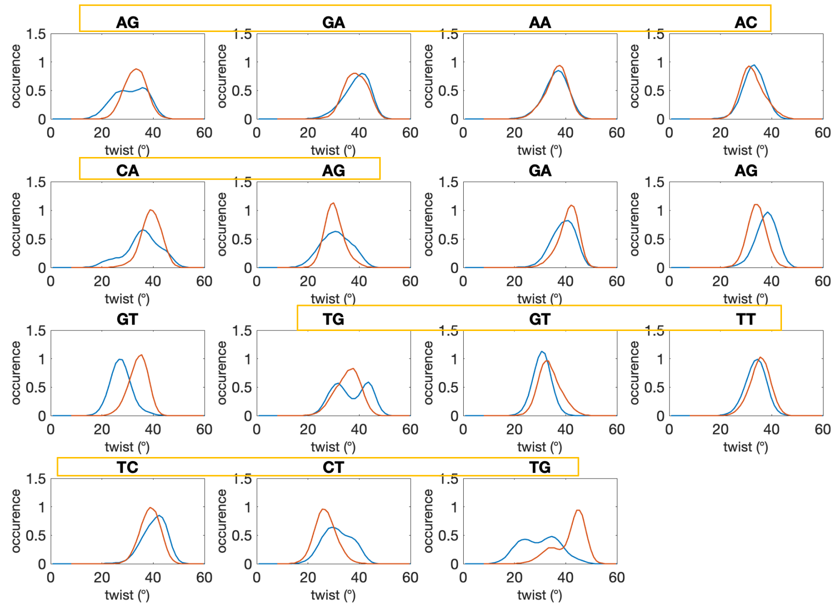
AG
GA
AA
AC
CA
AG
GA
AG
GT
TG
GT
TT
TC
CT
TG
occurence
twist (°)

**B**

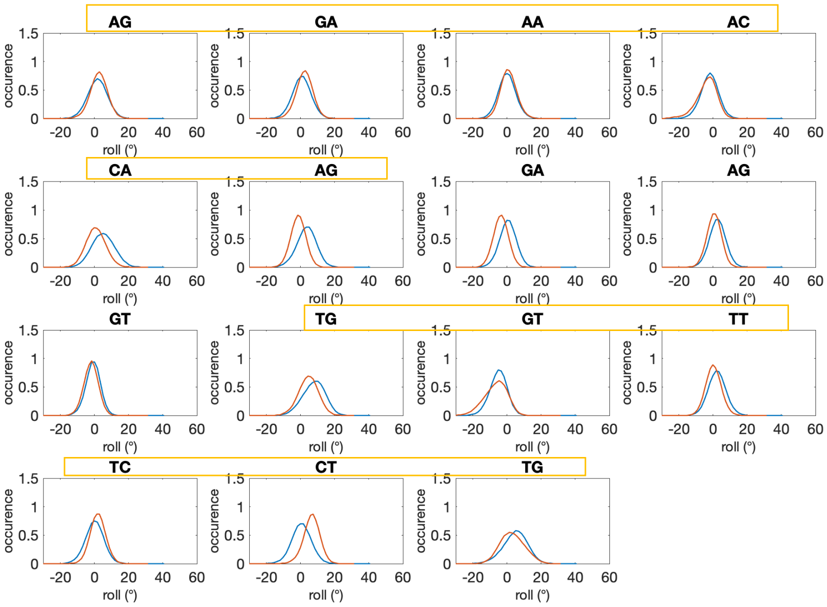
AG
GA
AA
AC
CA
AG
GA
AG
GT
TG
GT
TT
TC
CT
TG
occurence
roll (°)

**C**

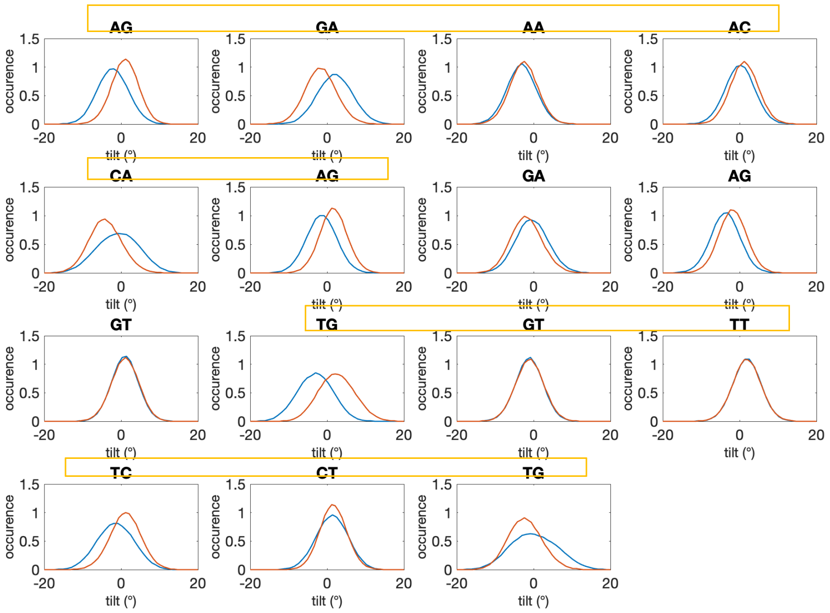

AG
GA
AA
AC
CA
AG
GA
AG
GT
TG
GT
TT
TC
CT
TG
occurence
tilt (°)


**D**

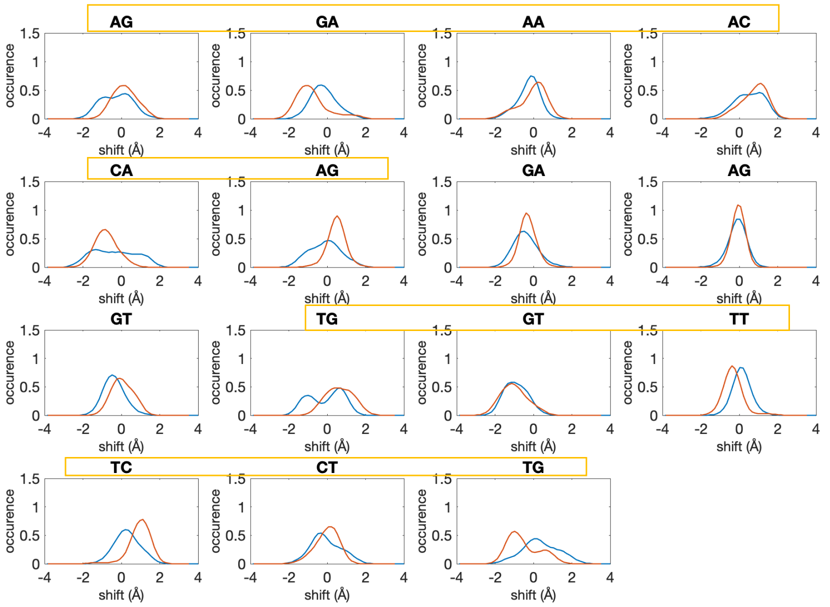

AG
GA
AA
AC
CA
AG
GA
AG
GT
TG
GT
TT
TC
CT
TG
occurence
shift (Å)

**E**

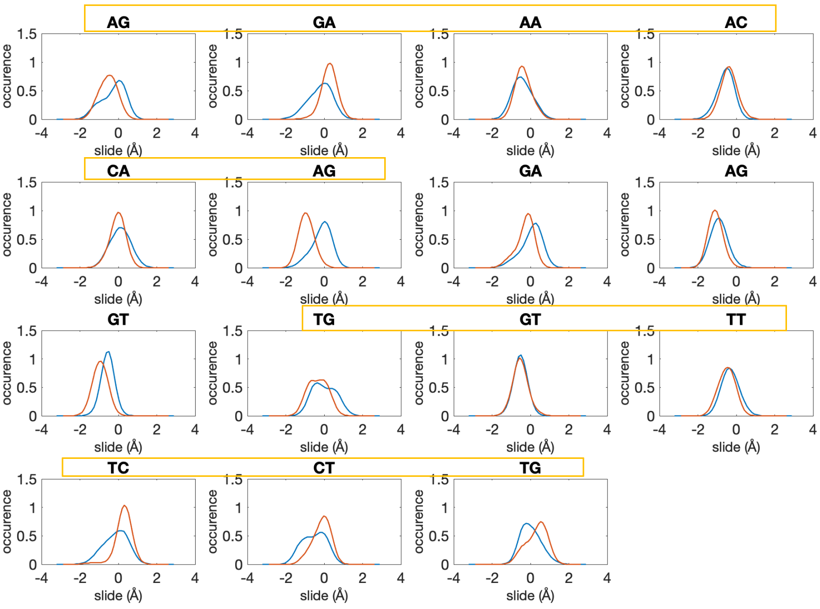


**F**

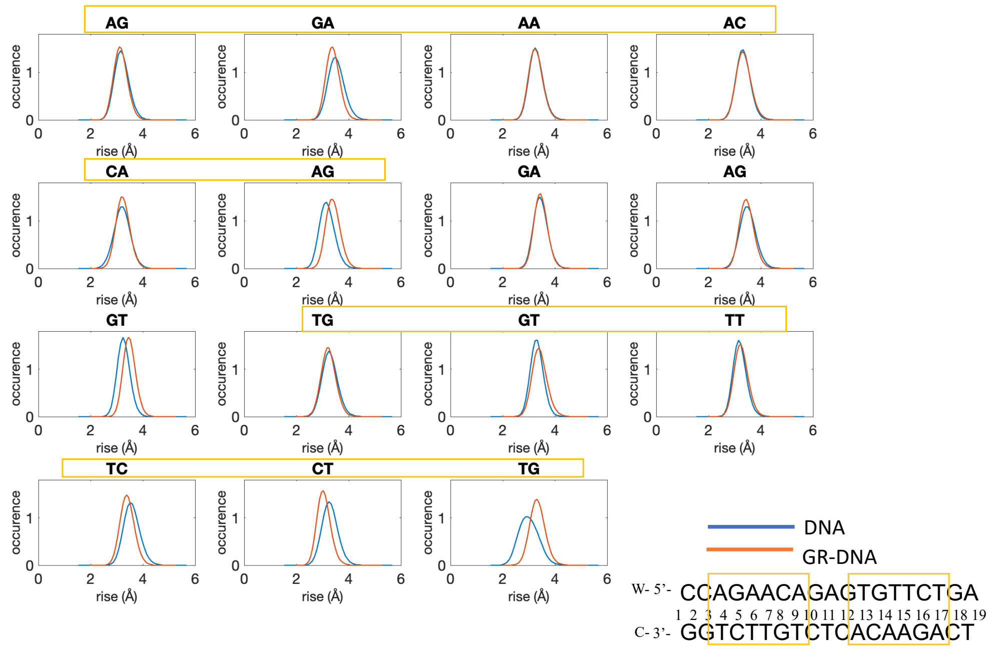


**Figure S11:** Helical parameters distributions for unbound and GR-bound DNA. **A.** Twist **B.** Roll **C.** Tilt **D.** Shift E. Slide **F.** Rise. The glucocorticoid receptor response elements 1 and 2 are highlighted with yellow.

**A**

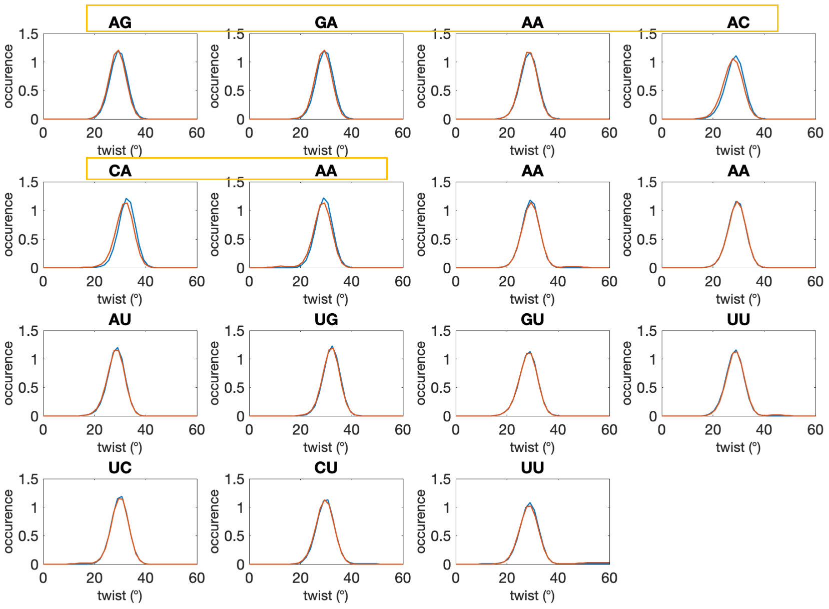
AG
GA
AA
AC
CA
AA
AA
AA
AU
UG
GU
UU
UC
CU
UU
occurence
twist (°)
1.5
1
0.5
0
0
20
40
60

**B**

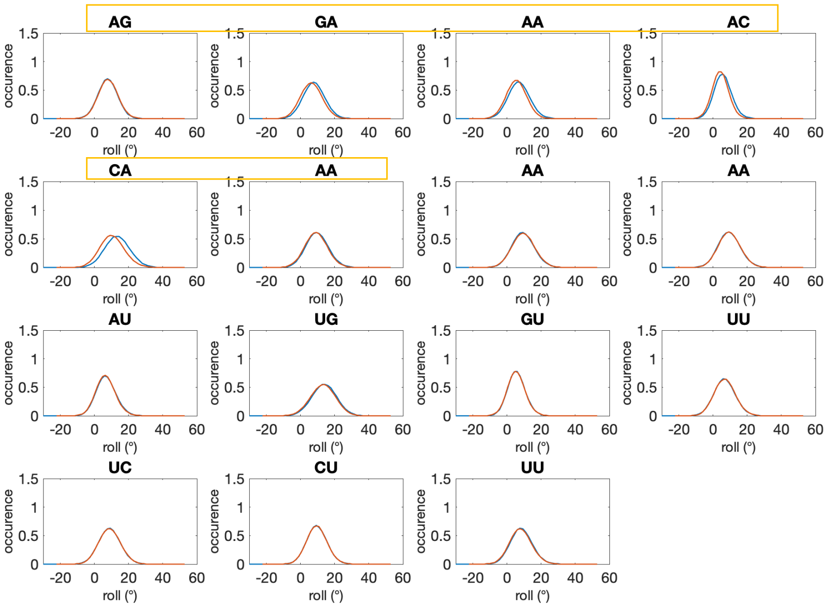
AG
GA
AA
AC
CA
AA
AA
AA
AU
UG
GU
UU
UC
CU
UU
occurence
roll (°)
1.5
1
0.5
0
-20
0
20
40
60

C

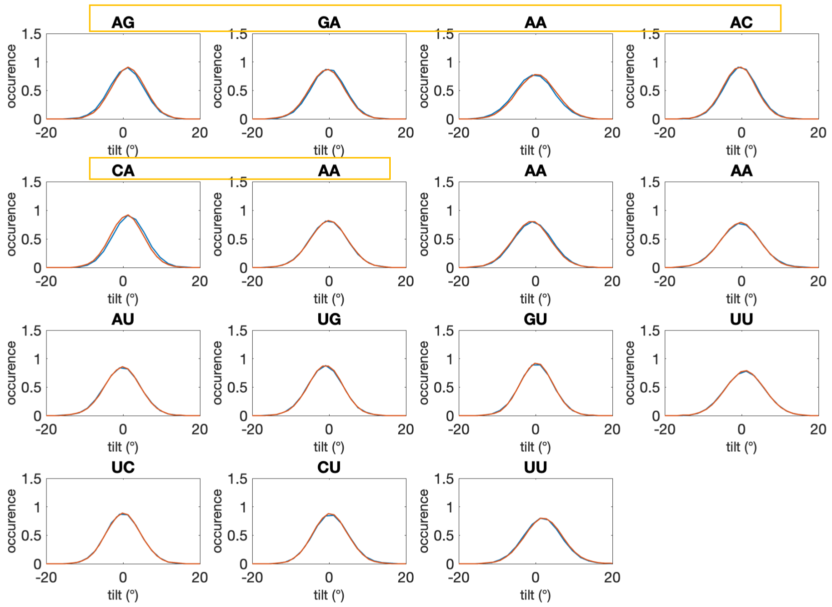
AG
GA
AA
AC
CA
AA
AA
AA
AU
UG
GU
UU
UC
CU
UU
occurence
tilt (°)

D

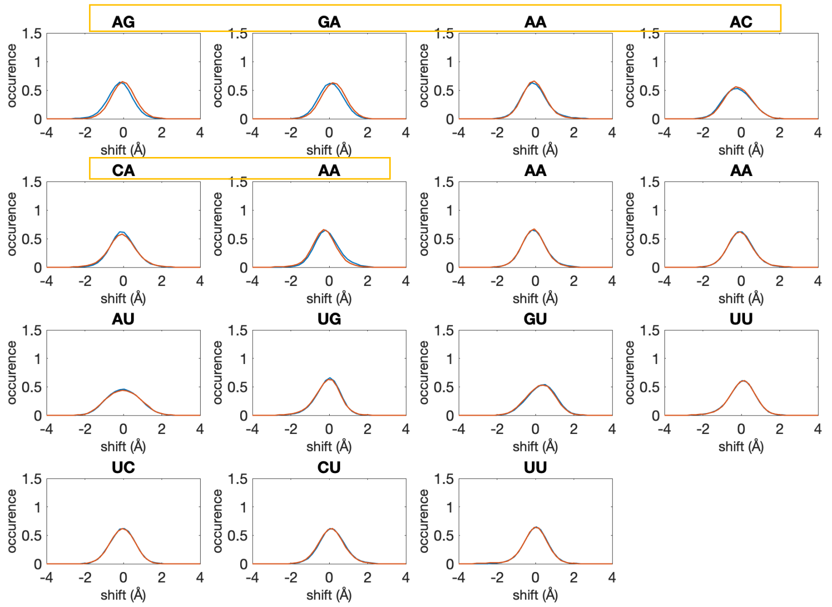
AG
GA
AA
AC
CA
AA
AA
AA
AU
UG
GU
UU
UC
CU
UU
occurence
shift (Å)

E

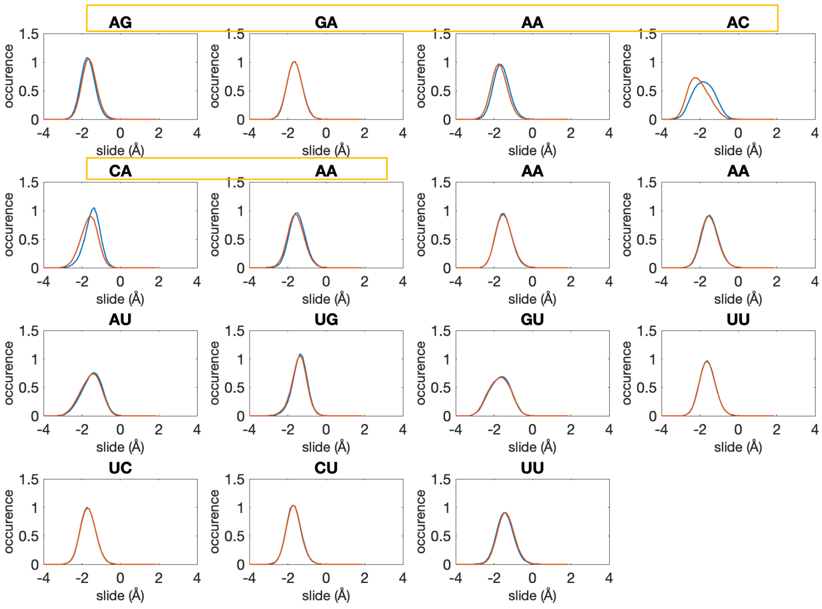


F

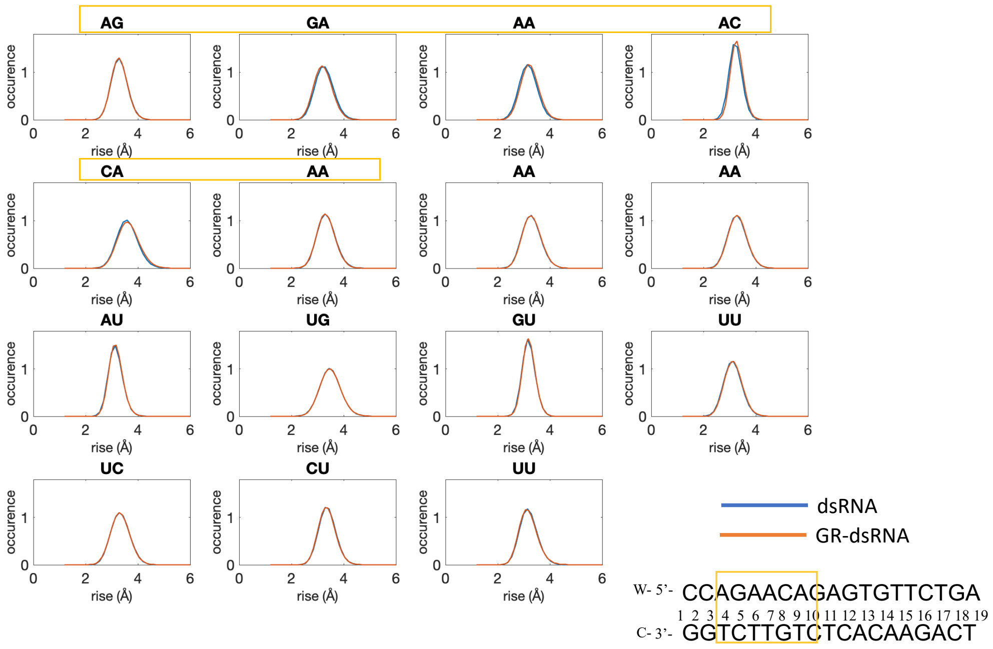


**Figure S12:** Helical parameters distributions for unbound and GR-bound dsRNA. **A.** Twist **B.** Roll **C.** Tilt **D.** Shift E. Slide **F.** Rise. The glucocorticoid receptor response element is highlighted with yellow.

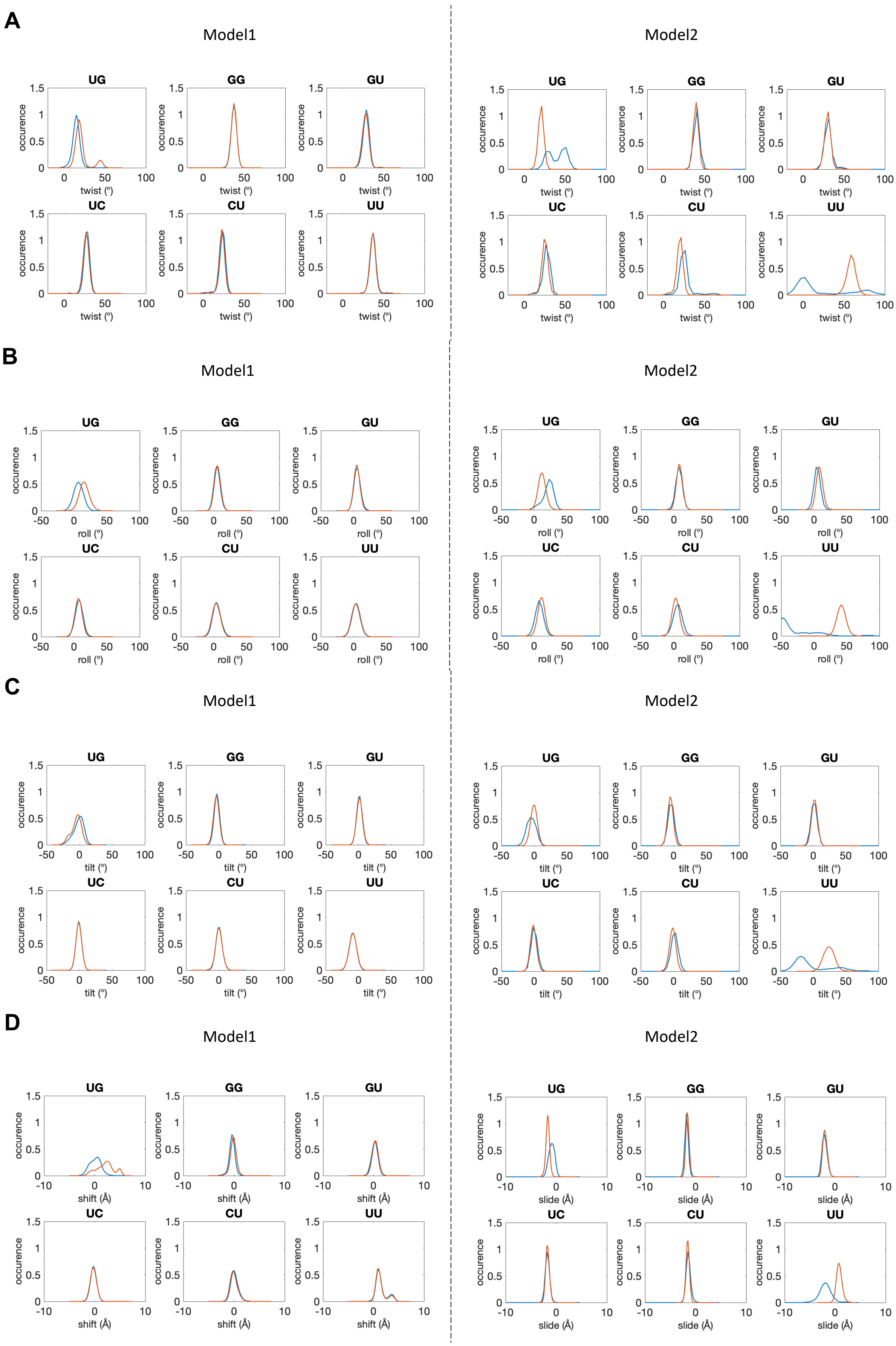
A
Model1
Model2
UG
GG
GU
UC
CU
UU
occurence
twist (°)
B
Model1
Model2
UG
GG
GU
UC
CU
UU
occurence
roll (°)
C
Model1
Model2
UG
GG
GU
UC
CU
UU
occurence
tilt (°)
D
Model1
Model2
UG
GG
GU
UC
CU
UU
occurence
shift (Å)
slide (Å)

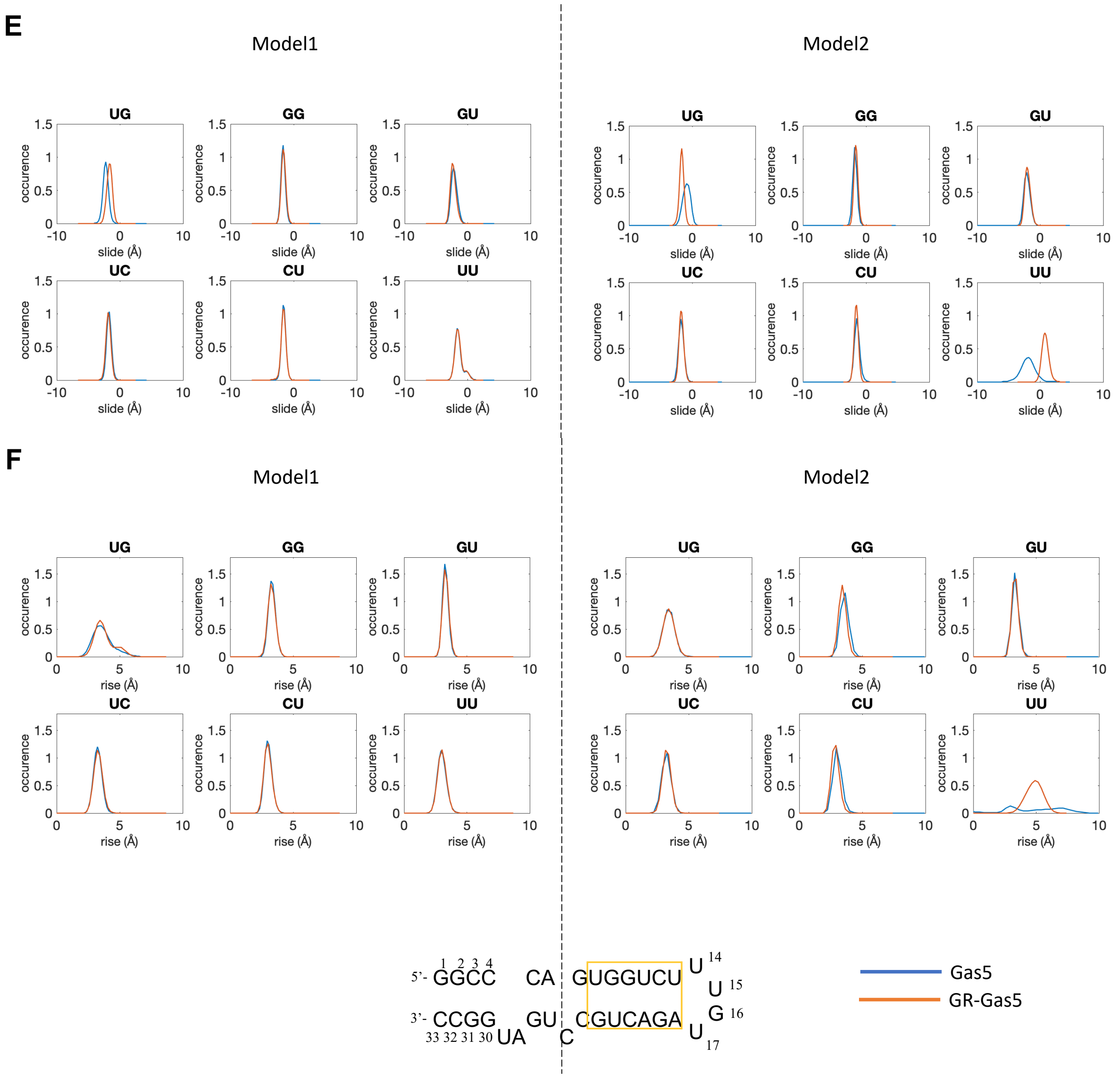


**Figure S13:** Helical parameters distributions of the glucocorticoid receptor response element for unbound and GR-bound Gas5_RNA for model1 (left panels) and model2 (right panels) . **A.** Twist **B.** Roll **C.** Tilt **D.** Shift E. Slide **F.** Rise.

**A**

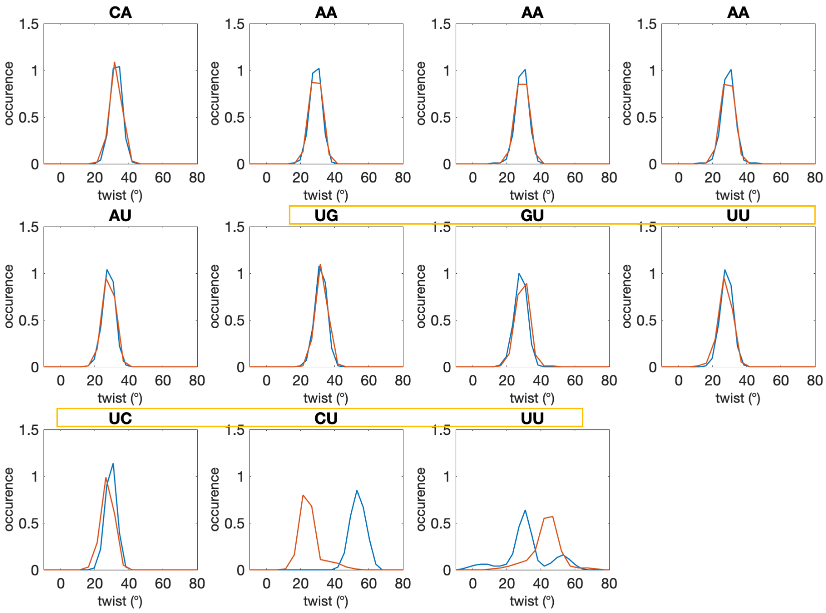
CA
AA
AA
AA
AU
UG
GU
UU
UC
CU
UU
occurence
twist (°)
0
0.5
1
1.5
0
20
40
60
80

**B**

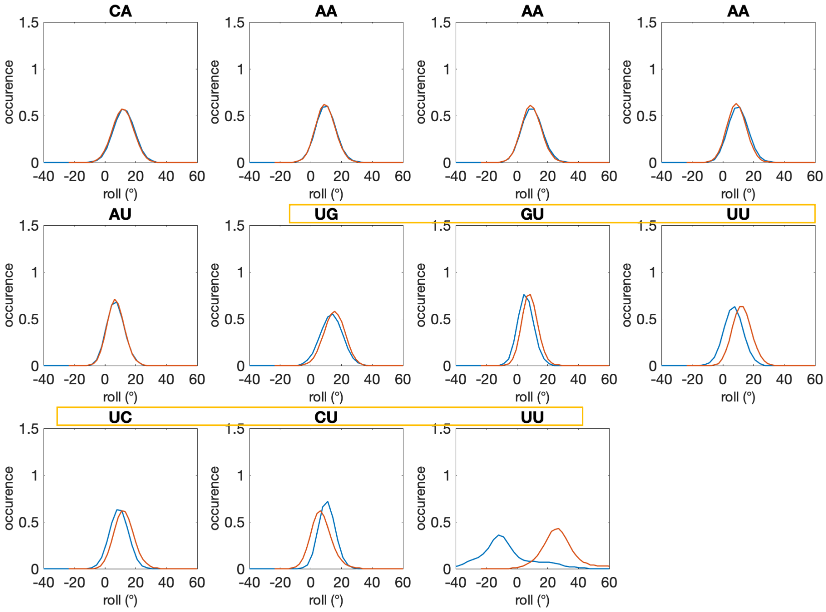
CA
AA
AA
AA
AU
UG
GU
UU
UC
CU
UU
occurence
roll (°)
0
0.5
1
1.5
-40
-20
0
20
40
60

**C**

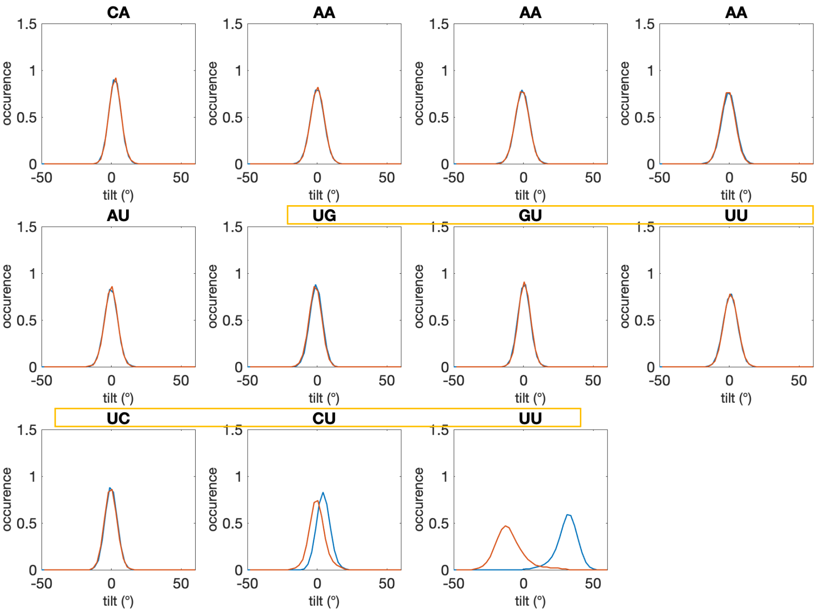
CA
AA
AA
AA
AU
UG
GU
UU
UC
CU
UU
occurence
tilt (°)
1.5
1
0.5
0
-50
0
50

**D**

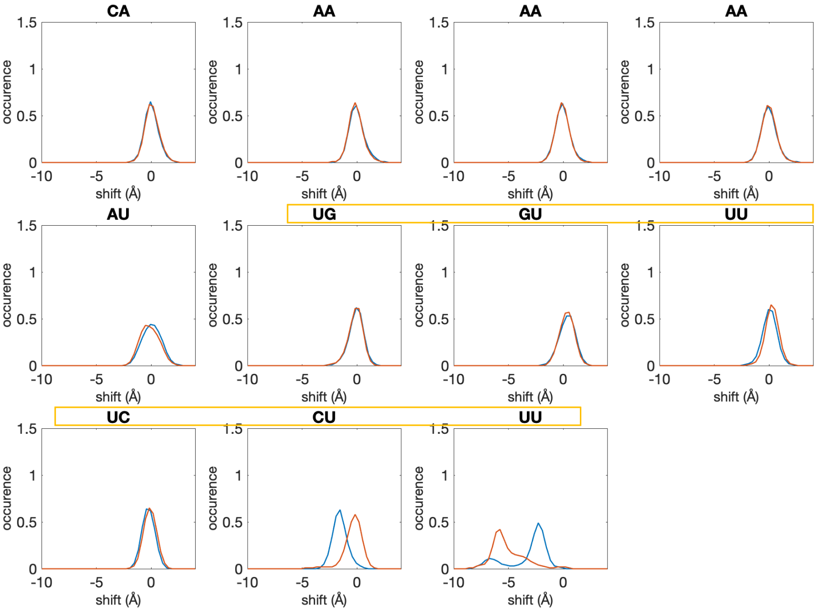
CA
AA
AA
AA
AU
UG
GU
UU
UC
CU
UU
occurence
shift (Å)
1.5
1
0.5
0
-10
-5
0

E

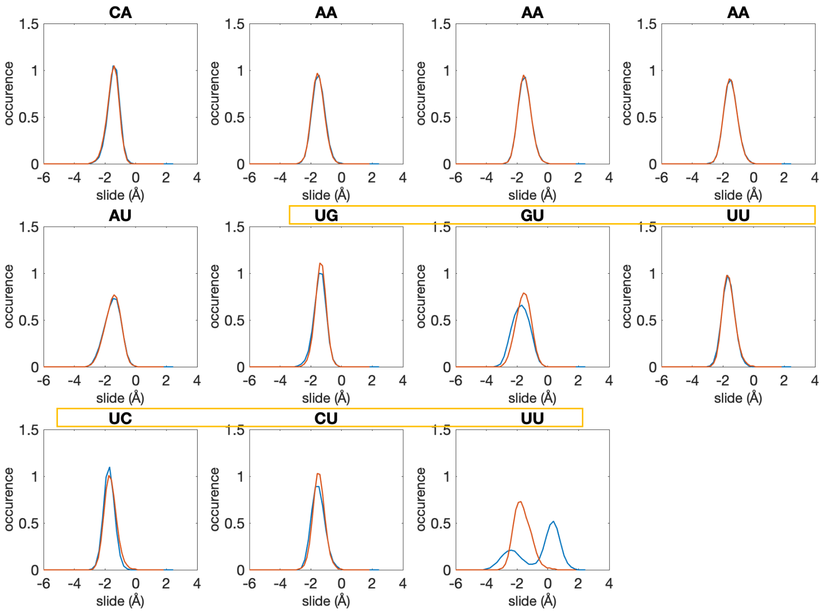


F

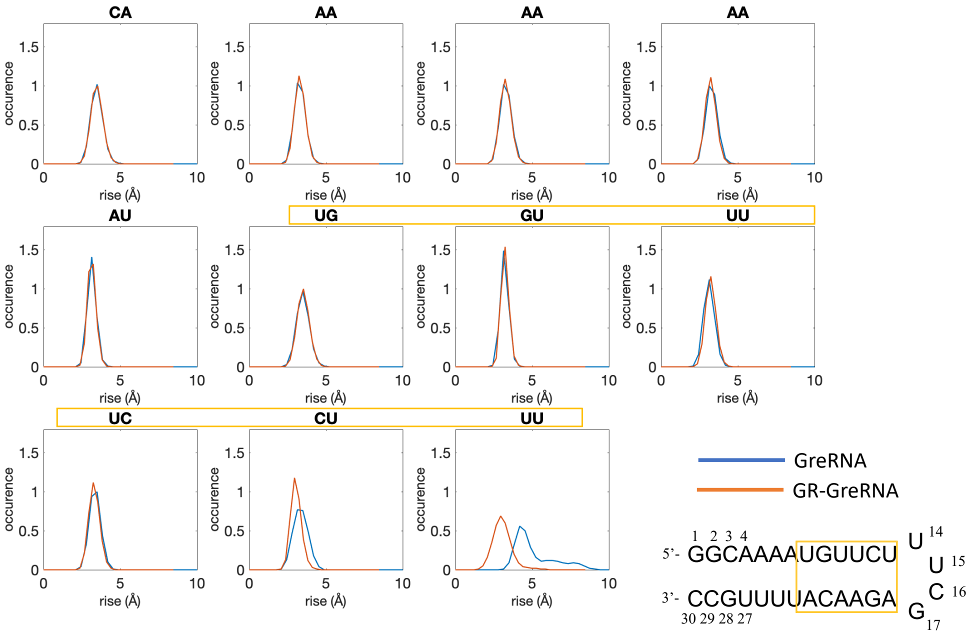


**Figure S14:** Helical parameters distributions for unbound and GR-bound GRE_RNA. **A.** Twist **B.** Roll **C.** Tilt **D.** Shift E. Slide **F.** Rise. The glucocorticoid receptor response element is highlighted with yellow.

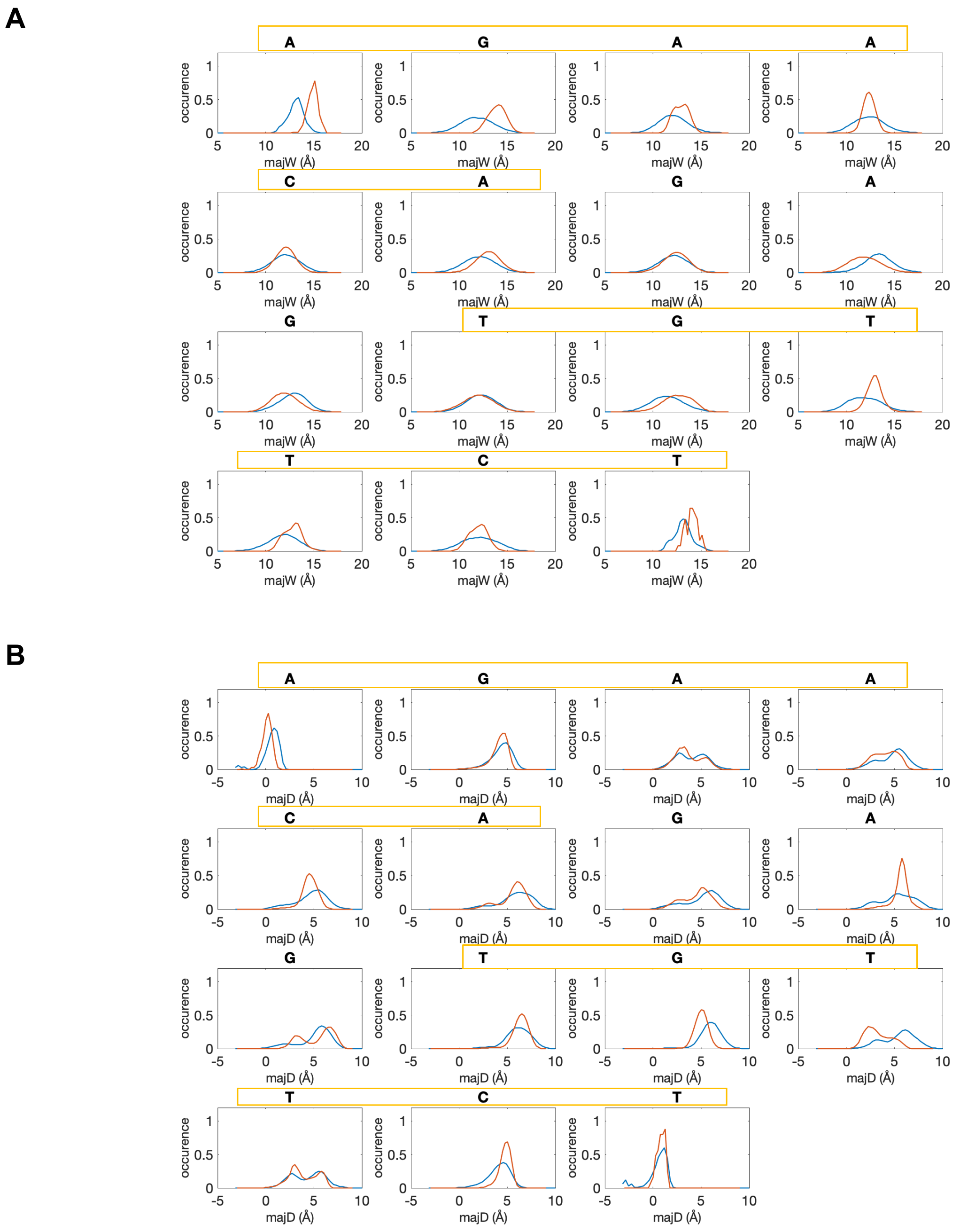
A
A G A A
C A G A
G T G T
T C T
occurence
majW (Å)
0 0.5 1
5 10 15 20
B
A G A A
C A G A
G T G T
T C T
occurence
majD (Å)
0 0.5 1
-5 0 5 10

C

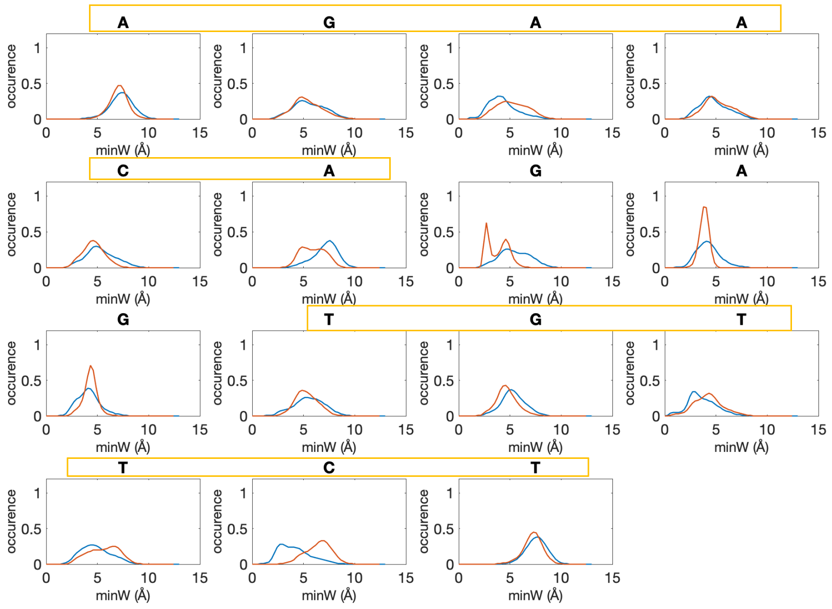


D

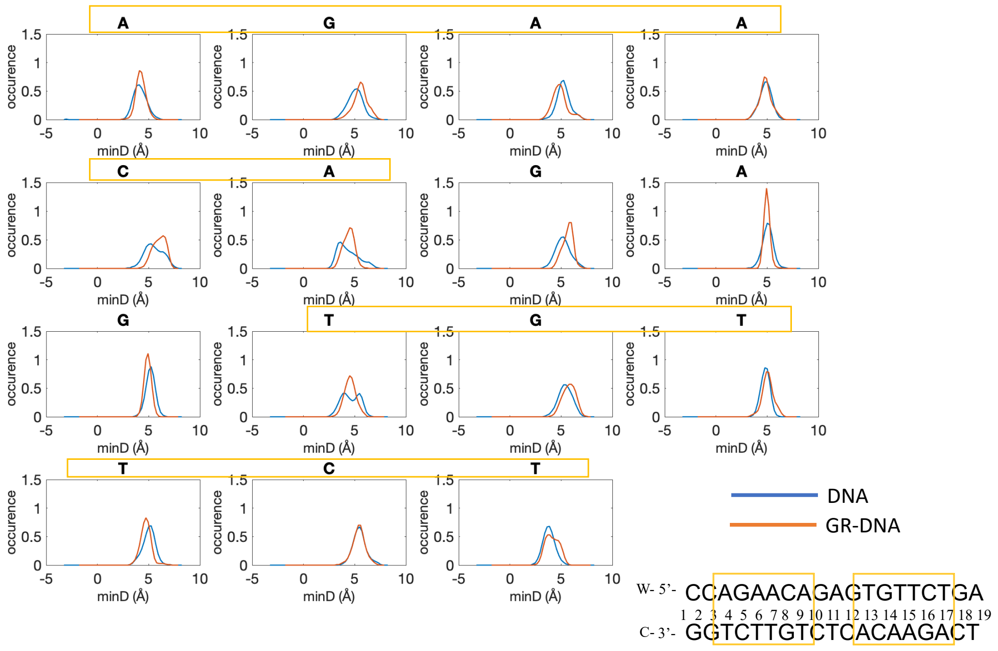


**Figure S15:** Groove parameters distributions for unbound and GR-bound DNA. **A.** Major groove width **B.** Major groove depth **C.** Minor groove width **D.** Minor groove depth. The glucocorticoid receptor response elements 1 and 2 are highlighted with yellow.

A

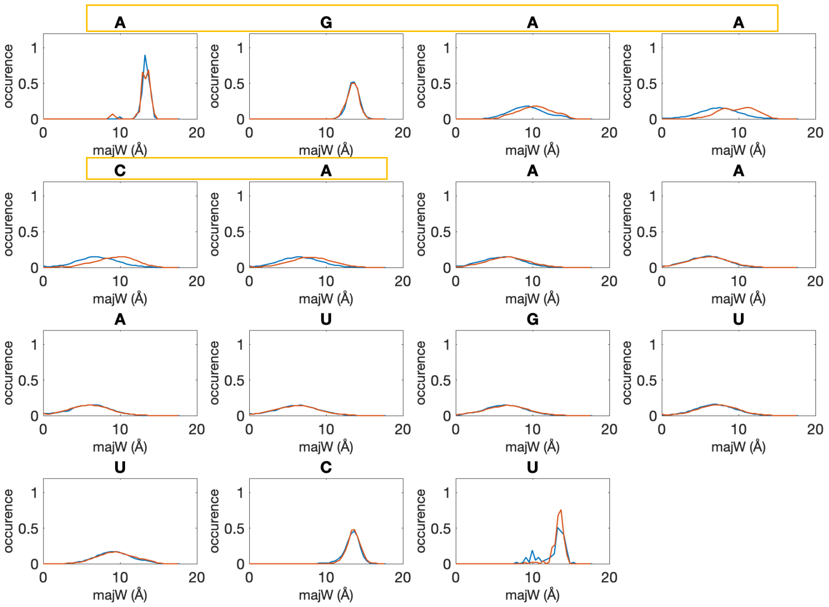
A
G
A
A
C
A
A
A
A
U
G
U
U
C
U
occurence
majW (Å)
0
0.5
1
10
20

B

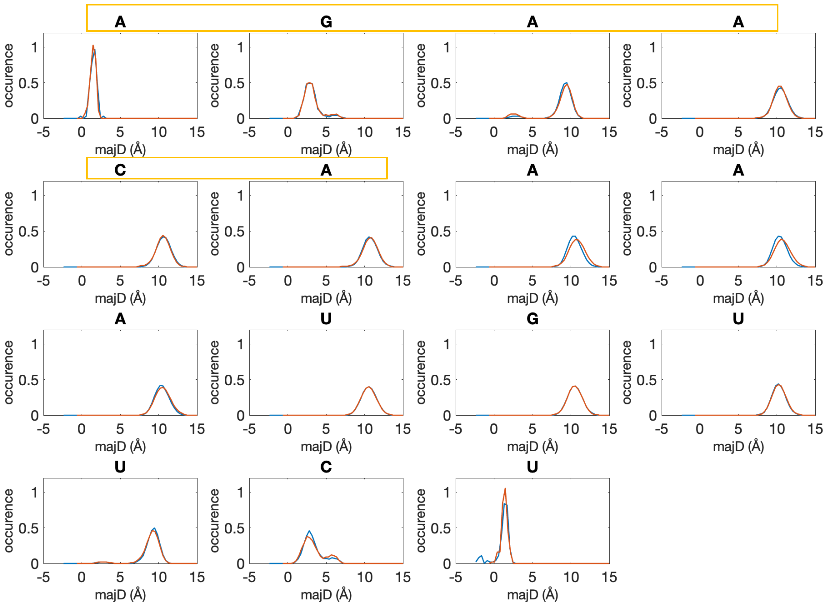
A
G
A
A
C
A
A
A
A
U
G
U
U
C
U
occurence
majD (Å)
0
0.5
1
-5
5
10
15

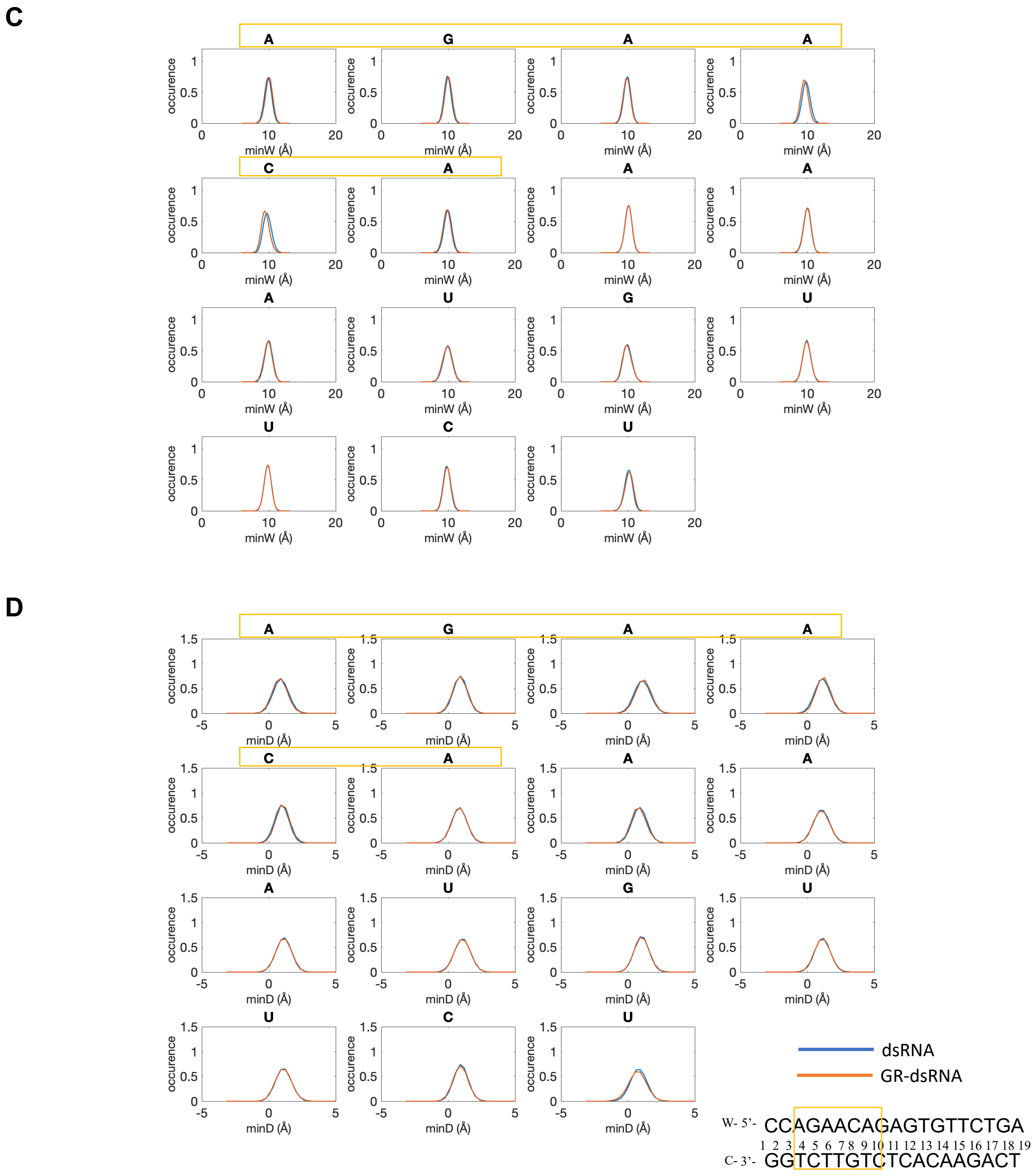


**Figure S16:** Groove parameters distributions for unbound and GR-bound dsRNA. **A.** Major groove width **B.** Major groove depth **C.** Minor groove width **D.** Minor groove depth. The glucocorticoid receptor response element is highlighted with yellow.

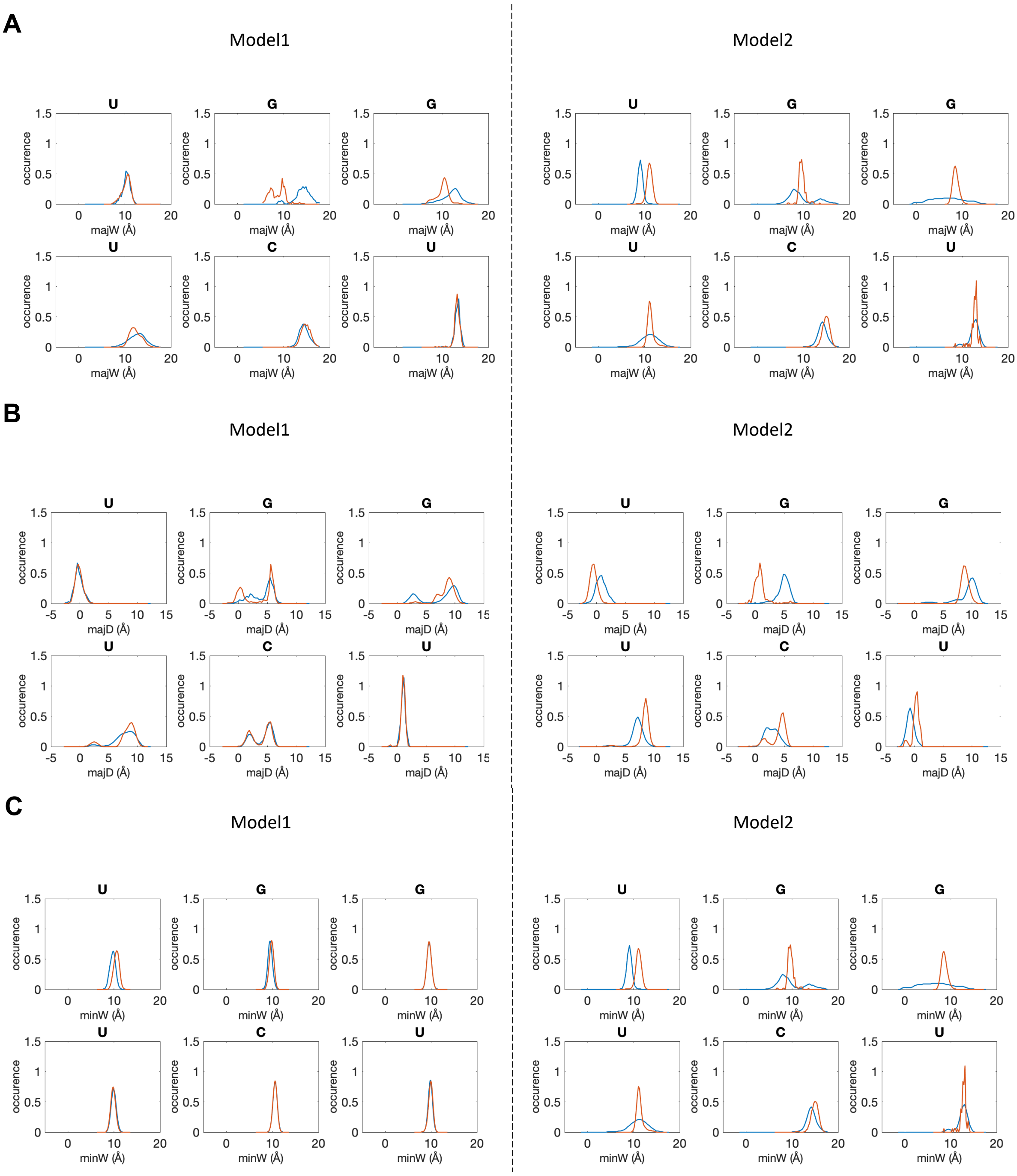
A
Model1
Model2
U
G
G
U
C
U
occurence
majW (Å)
B
Model1
Model2
majD (Å)
C
Model1
Model2
minW (Å)

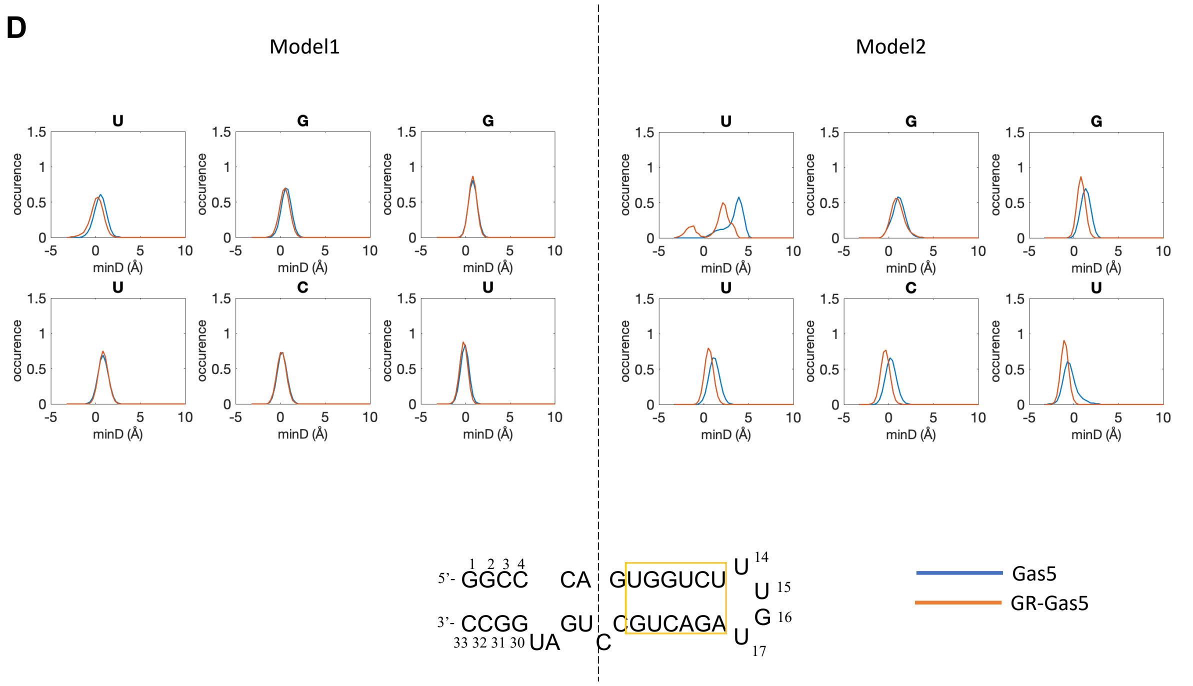


**Figure S17:** Groove parameters distributions of the glucocorticoid receptor response element for unbound and GR-bound Gas5_RNA. **A.** Major groove width **B.** Major groove depth **C.** Minor groove width **D.**

**A**

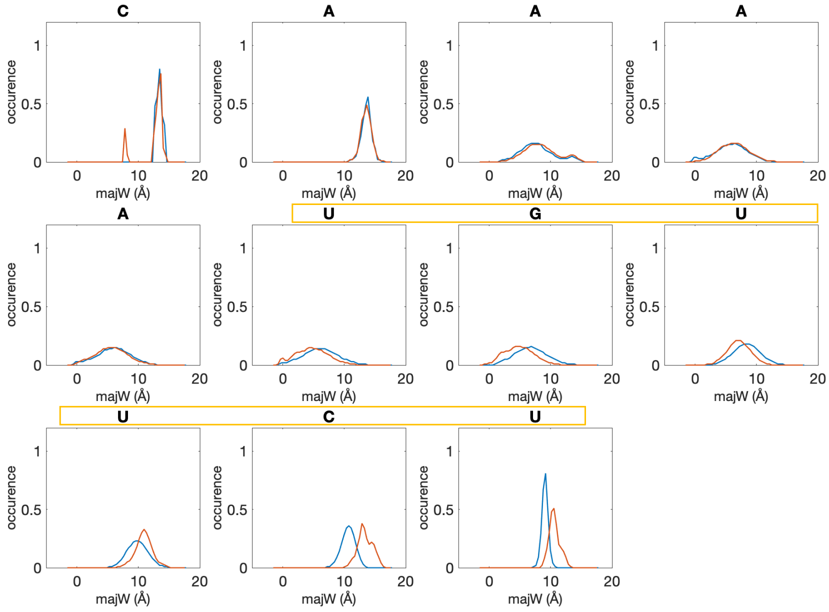
C
A
A
A
A
U
G
U
U
C
U
occurence
majW (Å)
0
0.5
1
10
20

**B**

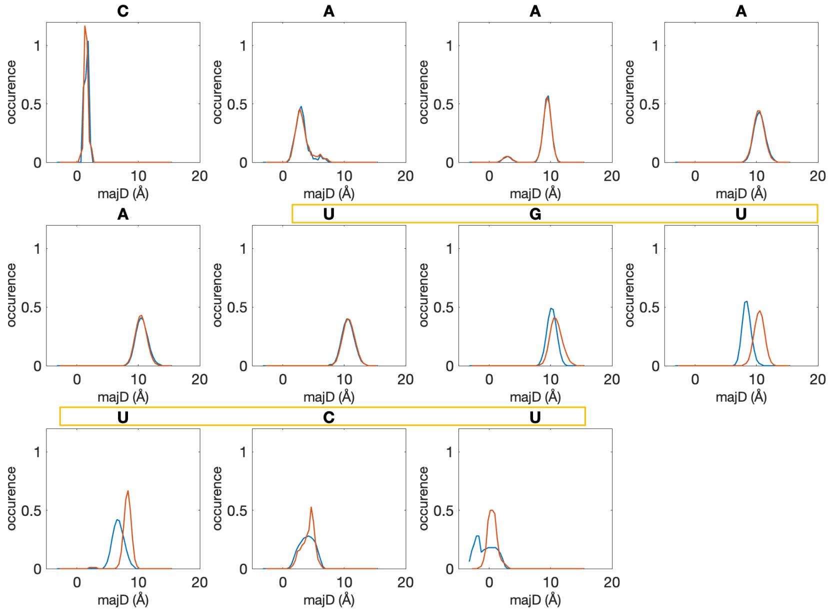
C
A
A
A
A
U
G
U
U
C
U
occurence
majD (Å)
0
0.5
1
10
20

**C**

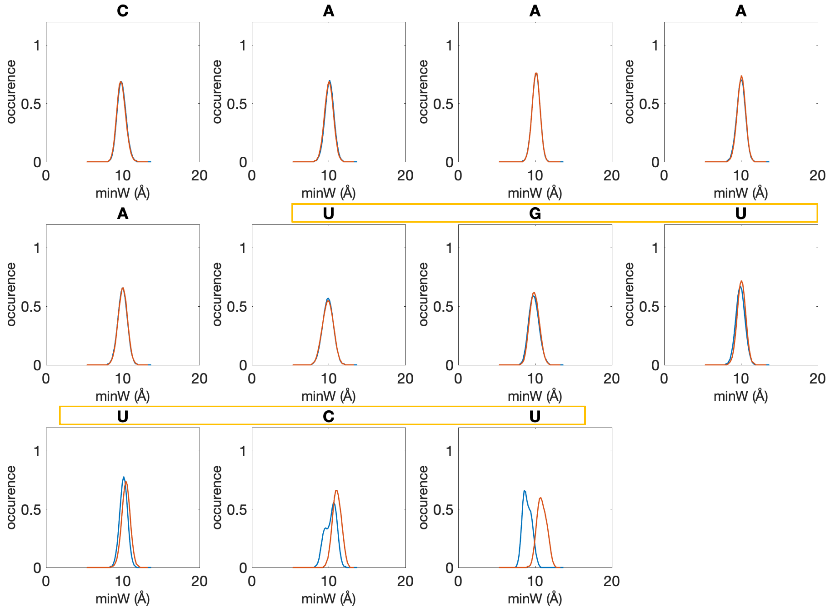


**D**

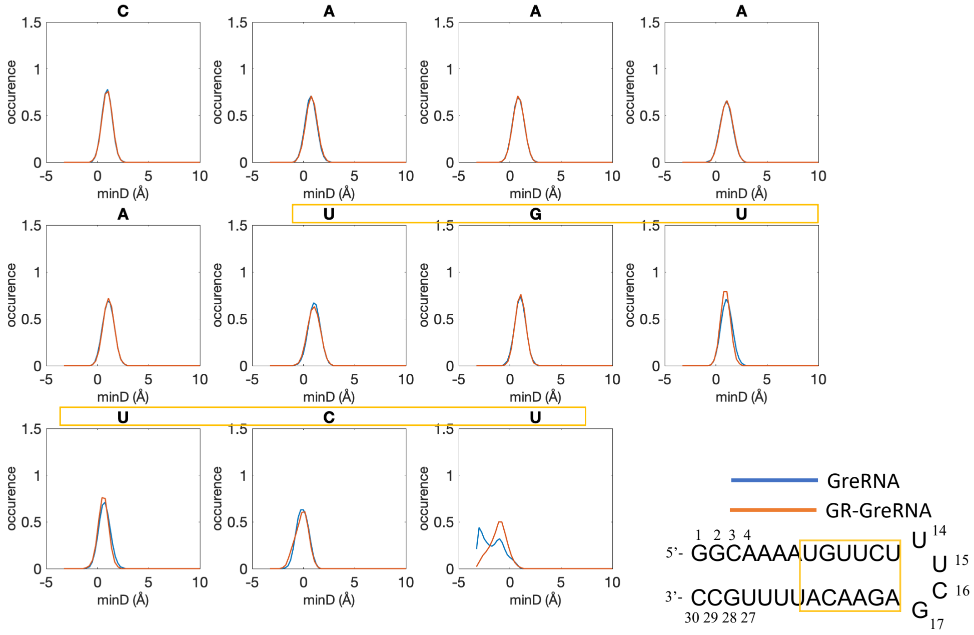


**Figure S18:** Groove parameters distributions for unbound and GR-bound GRE_RNA. **A.** Major groove width **B.** Major groove depth **C.** Minor groove width **D.** Minor groove depth. The glucocorticoid receptor response element is highlighted with yellow.

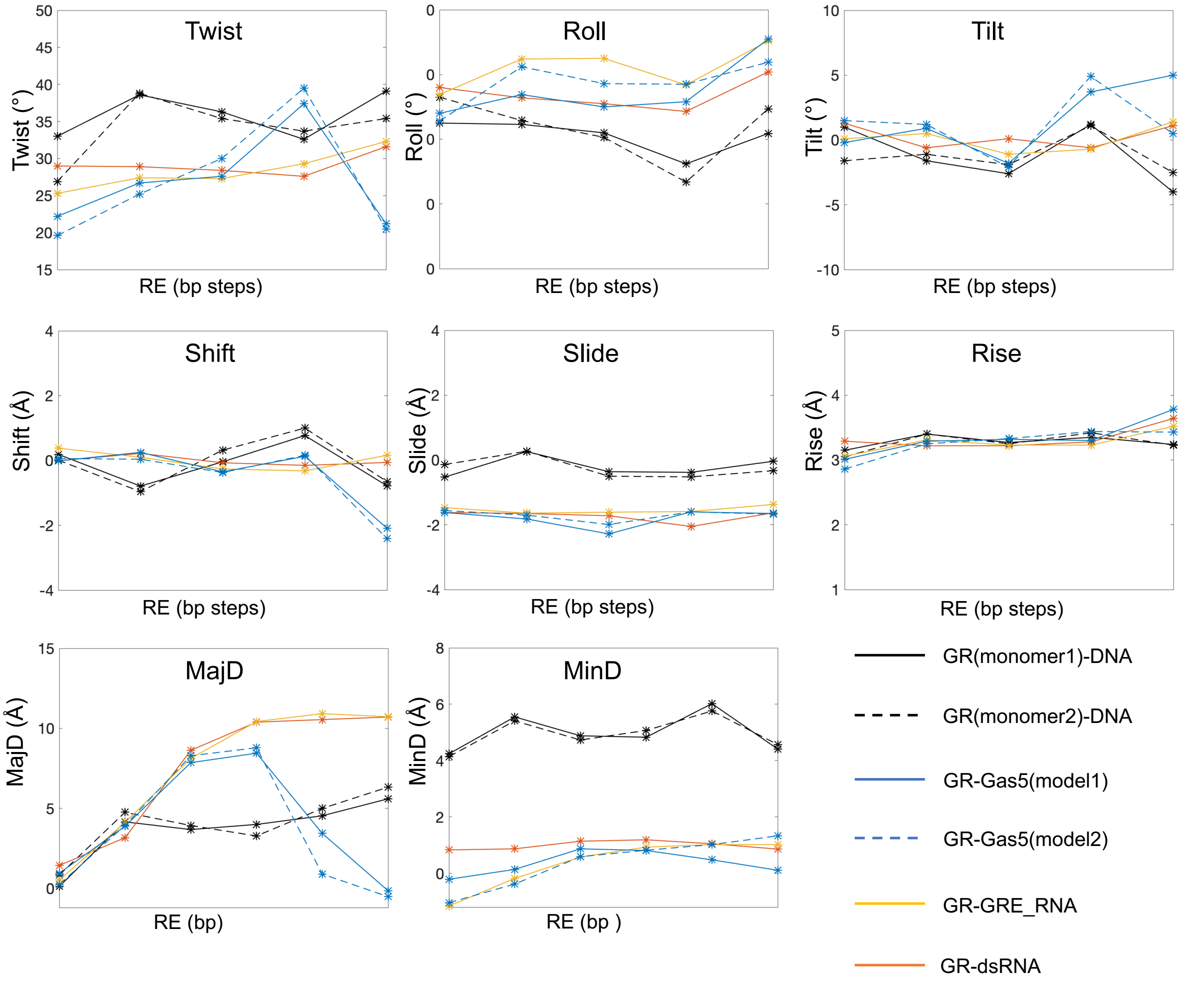


**Figure S19**: Comparison of average values for helical parameters and groove depths in different nucleic acids model systems for the b.p. within the GRE-site for the GR-bound systems.

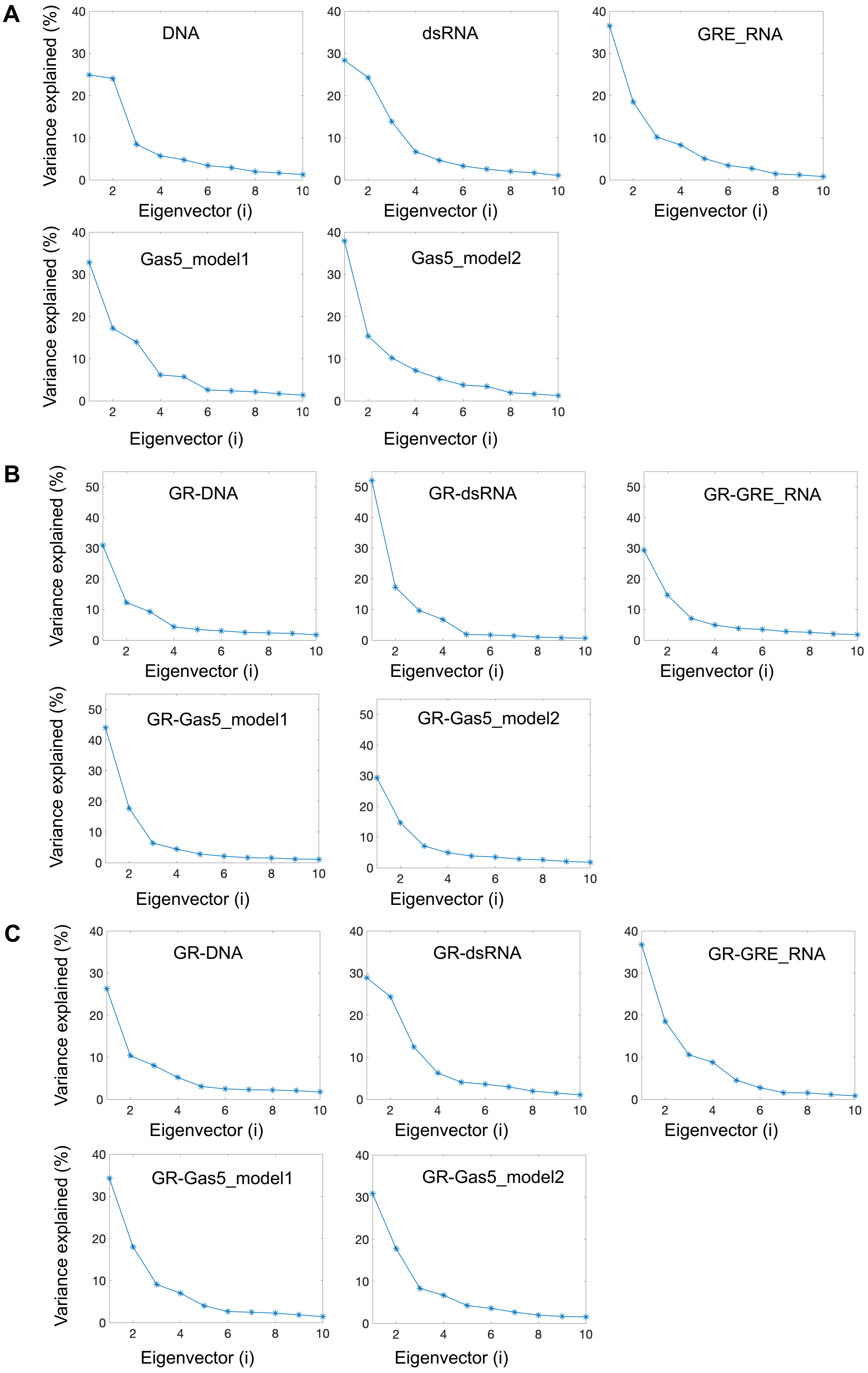
A
DNA
dsRNA
GRE_RNA
Gas5_model1
Gas5_model2
B
GR-DNA
GR-dsRNA
GR-GRE_RNA
GR-Gas5_model1
GR-Gas5_model2
C
GR-DNA
GR-dsRNA
GR-GRE_RNA
GR-Gas5_model1
GR-Gas5_model2
Variance explained (%)
Eigenvector (i)

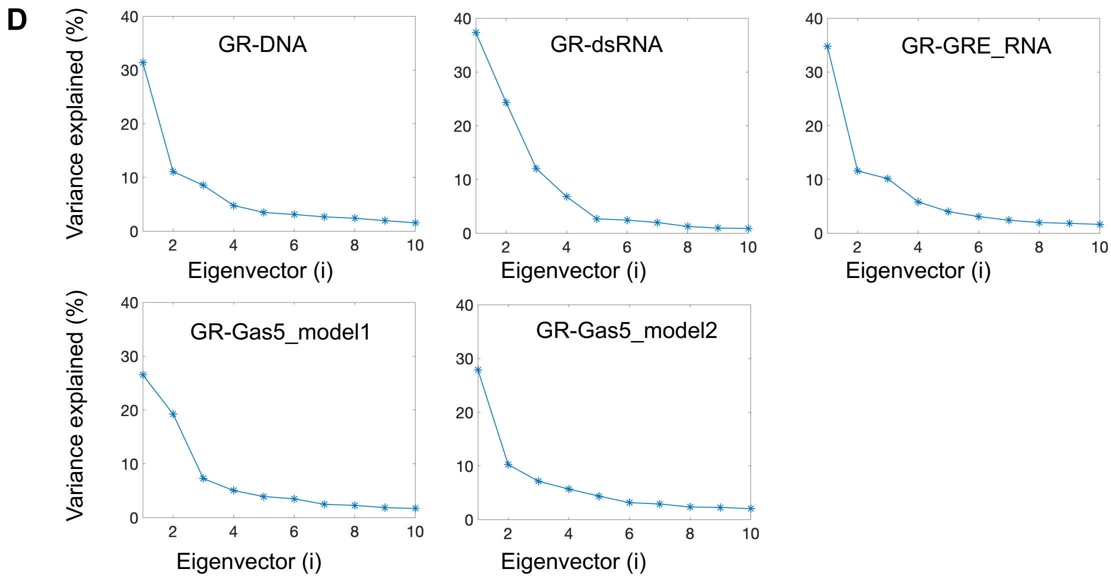


**Figure S20:** PCA analysis showing the variance explained by the first 10 eigenvectors. **A.** For unbound nucleic acids systems. For the GR-bound nucleic acids systems for the three PCA analyses performed: **B.** for the heavy atoms of the protein-DNA/RNA systems with the superposition on the nucleic acids and **C.** for heavy atoms of DNA/RNA with the superposition on the nucleic acids and **D.** for the heavy atoms of the complexes with superposition on the entire protein-DNA/RNA systems

**Table S1**: Variance of the dynamics explained by the first three eigenvectors

| | DNA | dsRNA | GRE_RNA | Gas5_model1 | Gas5_model2 |
|---|---|---|---|---|---|
| Unbound | 57.9% | 66.4% | 65.1% | 64.0% | 63.5% |
| GR-bound | | | | | |
| (1) | 52.5% | 79.0% | 58.5% | 68.2% | 51.1% |
| (2) | 44.7% | 65.7% | 65.8% | 61.3% | 56.9% |
| (3) | 51.0% | 74.7% | 56.5% | 53.0% | 45.3% |

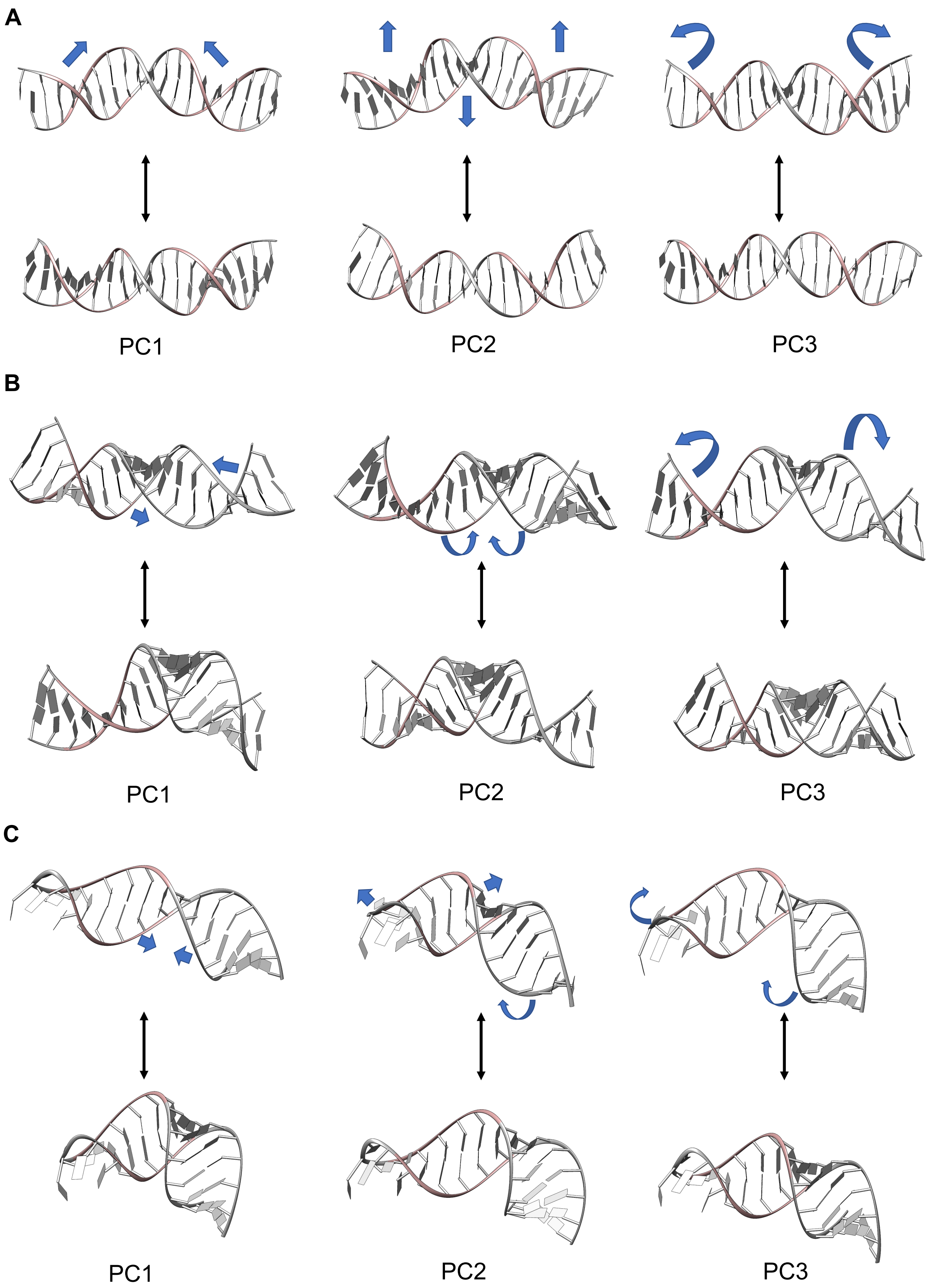
A
PC1
PC2
PC3
B
PC1
PC2
PC3
C
PC1
PC2
PC3

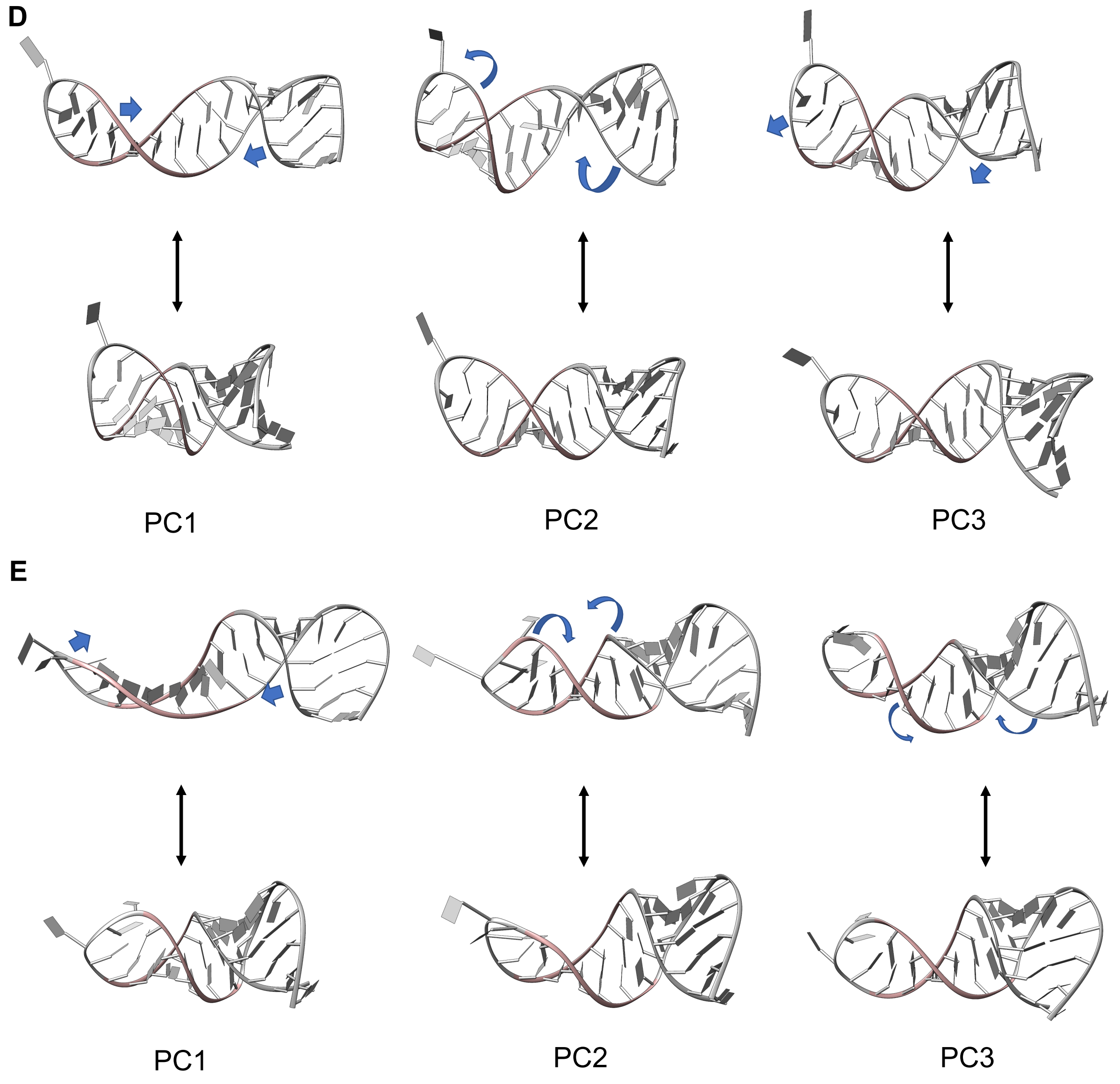


**Figure S21**: The extreme points for the first three PCs of the unbound nucleic acid systems. **A.** DNA **B.** dsRNA **C.** GRE_RNA **D.** Gas5_RNA_model1 **E.** Gas5_RNA_model2. The blue arrows indicate the motion described by the corresponding PC.

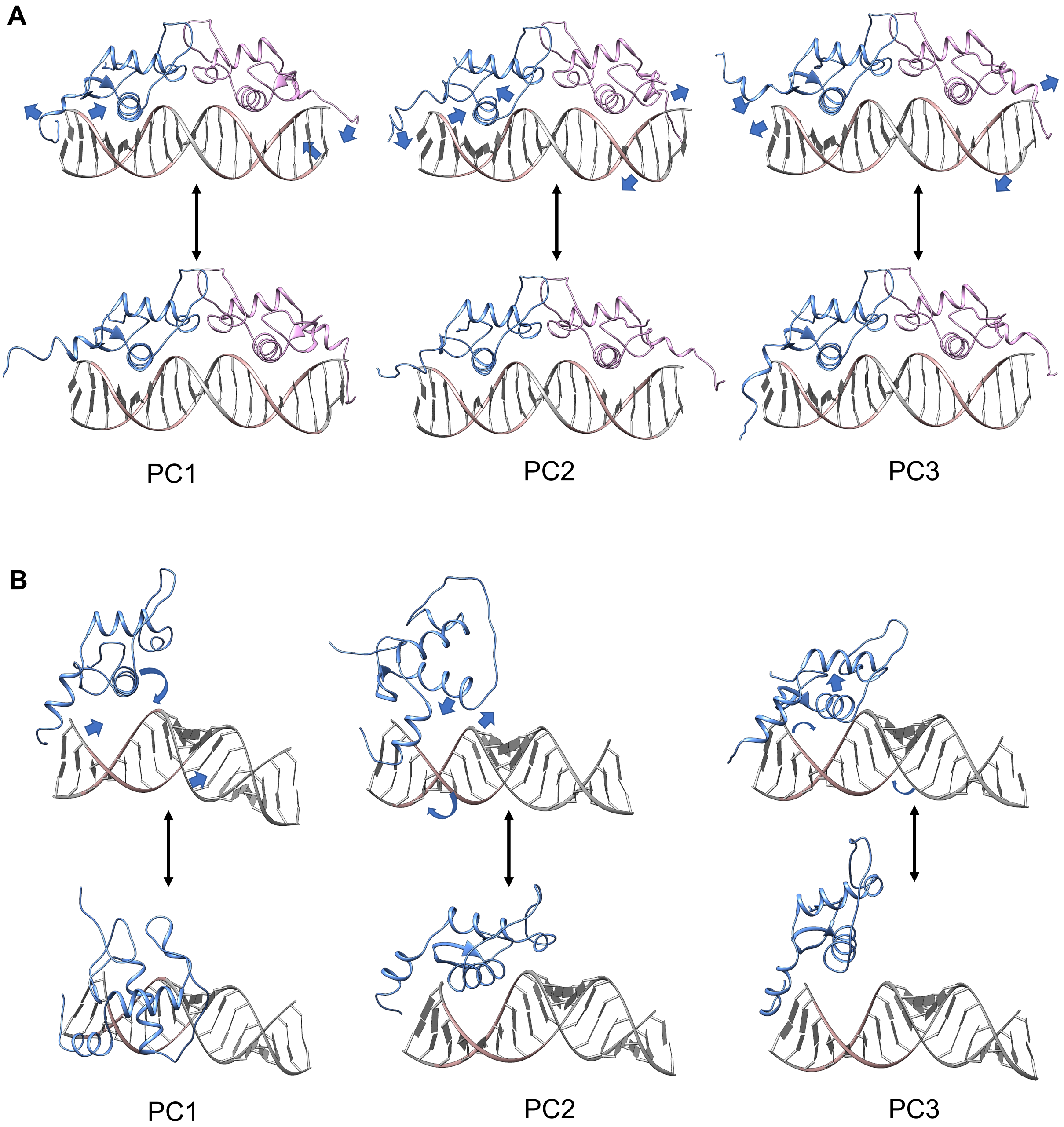
A
PC1
PC2
PC3
B
PC1
PC2
PC3

C

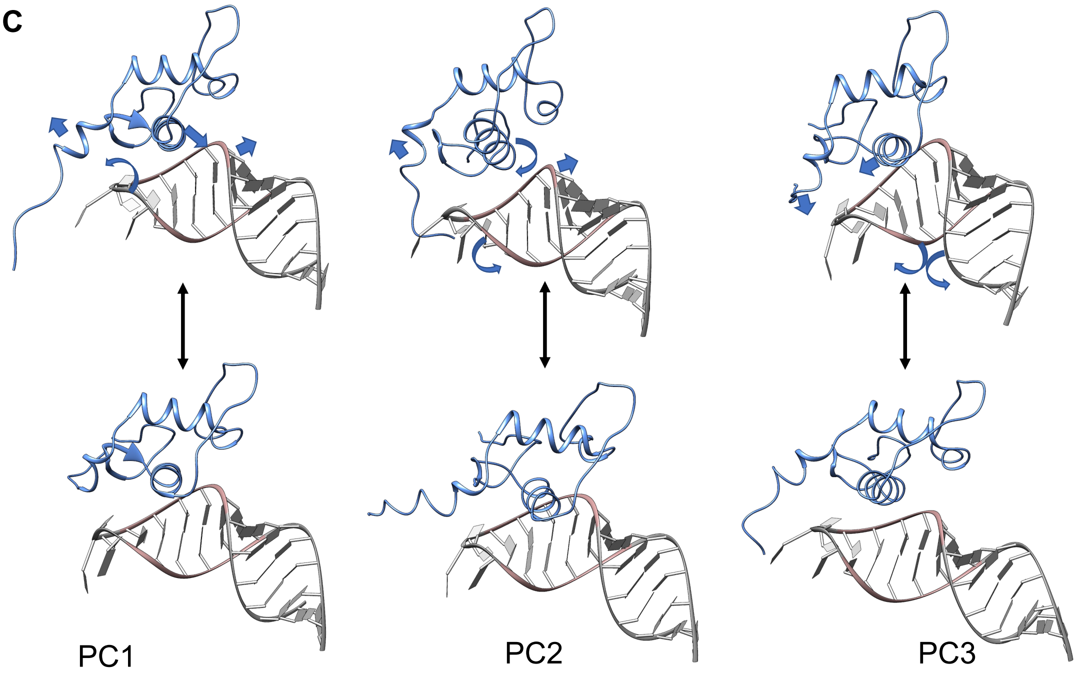


D

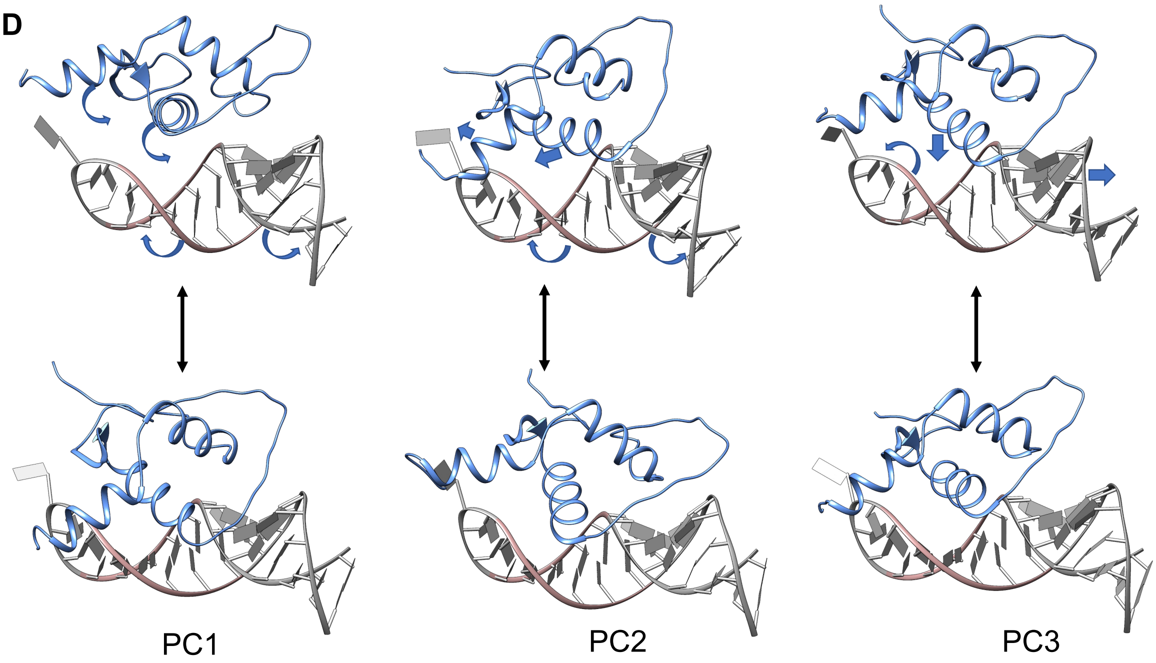

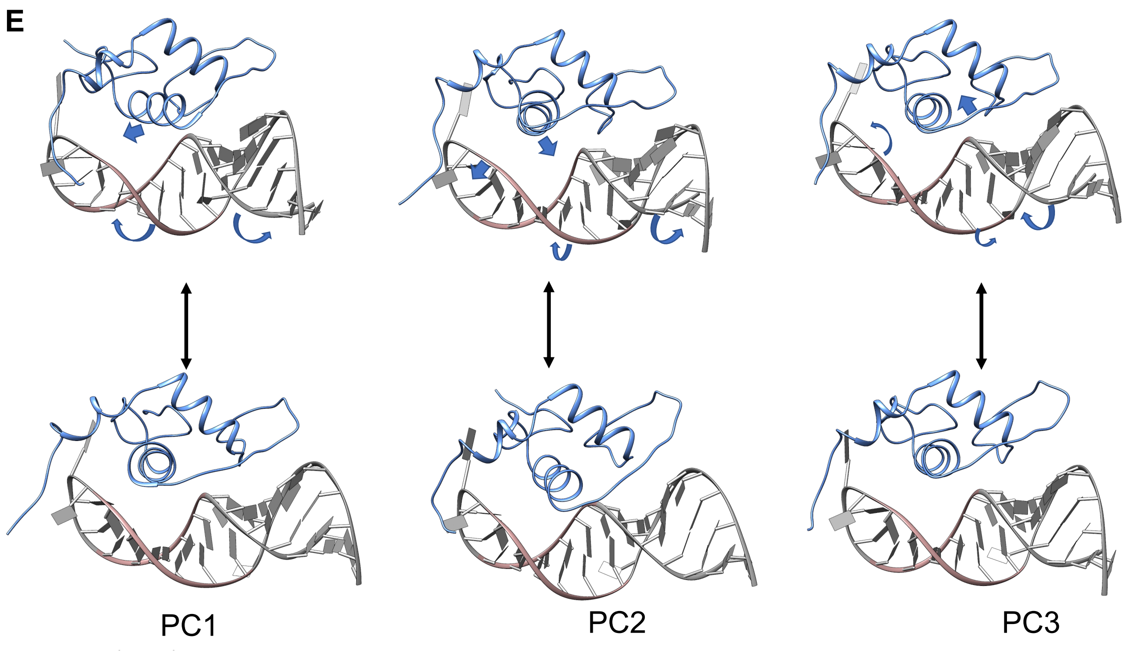


**Figure S22:** The extreme points for the first three PCs of the GR-bound **A.** DNA **B.** dsRNA **C.** GRE_RNA **D.** Gas5_RNA_model1 **E.** Gas5_RNA_model2. Representative motions for analysis (1) and (3). Where analysis (1) is performed for the heavy atoms of the protein-DNA/RNA systems with the superposition on the nucleic acids; and analysis (3) is performed for the heavy atoms of the protein-nucleic acids complexes with the superposition on the entire protein-DNA/RNA systems. The blue arrows indicate the motion described by the corresponding PC.

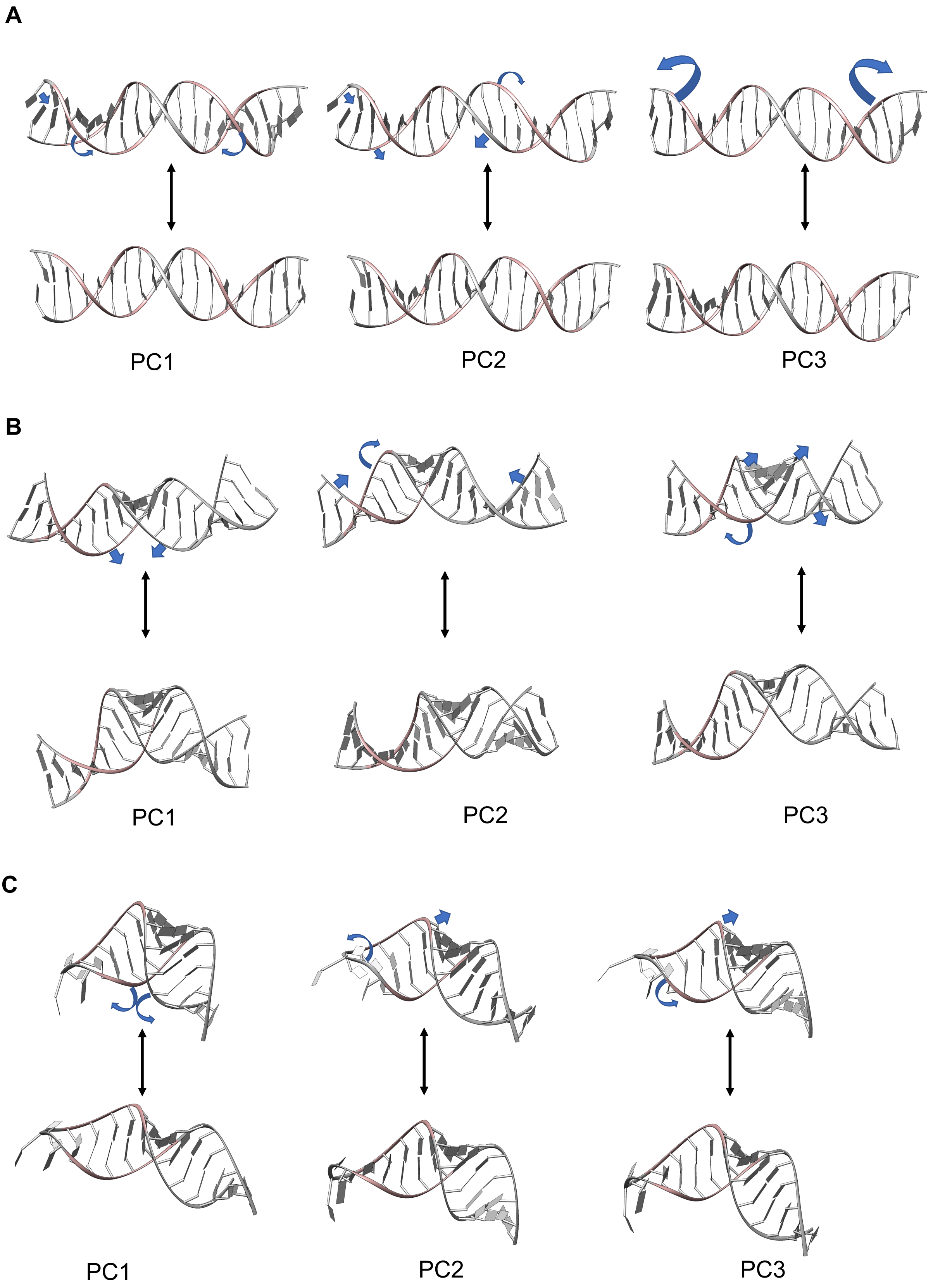
A
PC1
PC2
PC3
B
PC1
PC2
PC3
C
PC1
PC2
PC3

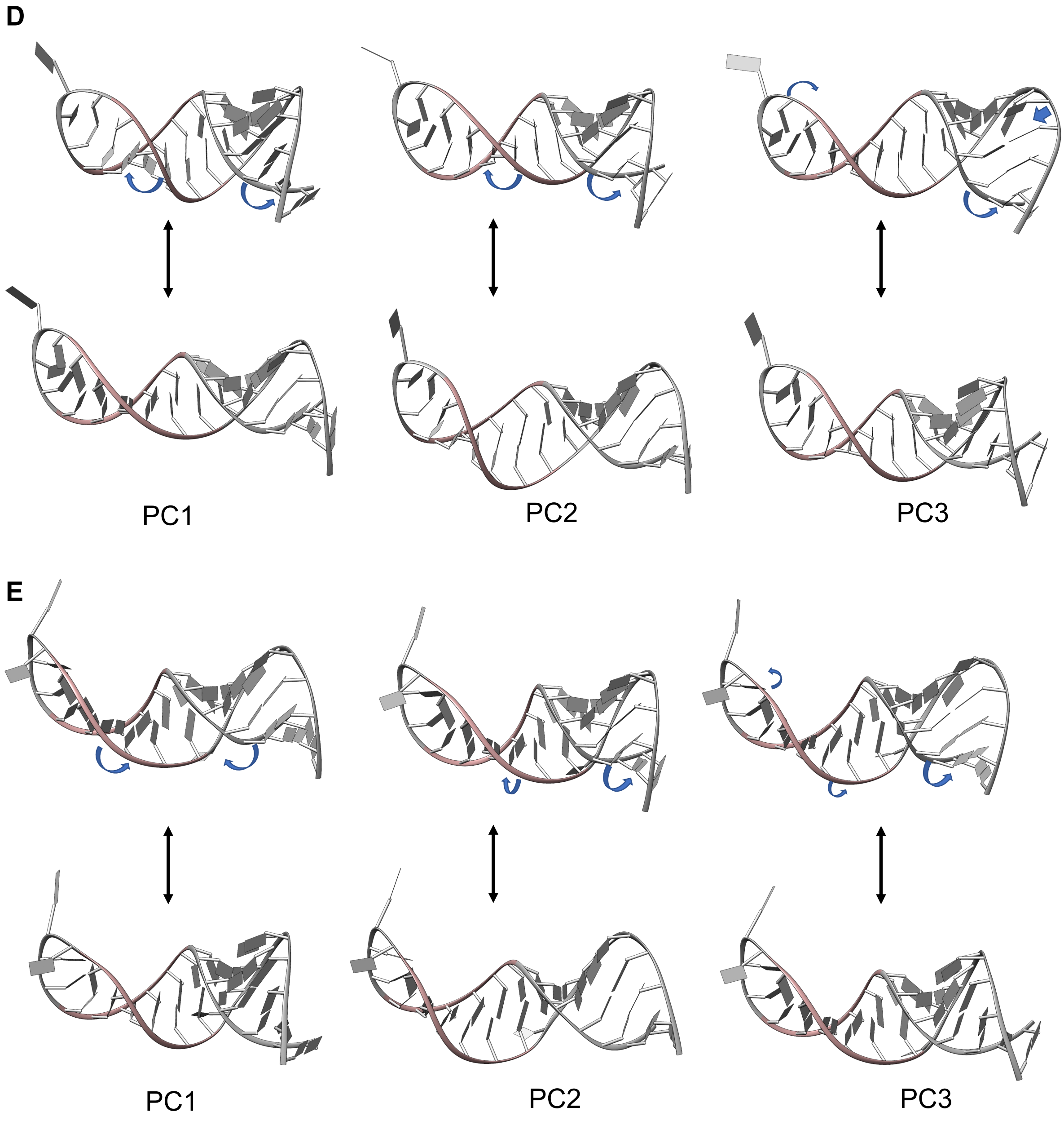


**Figure S23:** The extreme points for the first three PCs of the GR-bound **A.** DNA **B.** dsRNA **C.** GRE_RNA **D.** Gas5_RNA_model1 **E.** Gas5_RNA_model2. Representative motions for analysis (2), which is performed for heavy atoms of DNA/RNA with the superposition on the nucleic acids. The blue arrows indicate the motion described by the corresponding PC.
